# Stochasticity and heterogeneity restrict tipping points and alternative stable states

**DOI:** 10.64898/2026.06.15.732263

**Authors:** Santanu Das, Nadav Shnerb, Niv DeMalach

**Affiliations:** Department of Physics, Bar-Ilan University, Ramat-Gan IL52900, Israel; Institute of Plant Sciences and Genetics in Agriculture, Robert H. Smith Faculty of Agriculture, Food and Environment, Hebrew University of Jerusalem, Rehovot, Israel

## Abstract

Gradual environmental stress may trigger abrupt tipping points, trapping ecosystems in degraded states that are difficult to reverse. This possibility has strongly influenced ecosystem management, yet empirical evidence for such behavior remains mixed. A major unresolved question is how broadly stochasticity and heterogeneity restrict hysteresis, and whether their effects accumulate. To address it, we reconstructed full phase diagrams for four canonical ecological models (dryland desertification, grazing, insect outbreaks, and lake eutrophication) across gradients of stress, demographic stochasticity, environmental fluctuations, spatial heterogeneity, and dispersal. In every model, stochasticity and heterogeneity narrowed the region supporting alternative stable states and made the remaining transition less abrupt. Dispersal set their effective strength by averaging across space. These complexities eroded hysteresis along different routes -toward the degraded state, through a continuous transition, or, counterintuitively, toward the healthy state. Effects were largely additive, so several moderate complexities could eliminate hysteresis even when no single factor did. Tipping-point claims should therefore be tested under stochastic, heterogeneous conditions.

## INTRODUCTION

Ecosystems worldwide are under mounting pressure from human activity, including climate change, land-use transformation, and pollution [1–8]. Drylands face intensifying droughts and expanding desertification [9–11] while overgrazing threatens the productivity of rangelands that support billions of people [11, 12]. At the same time, nutrient pollution is degrading water quality and biodiversity across freshwater, marine, and terrestrial systems [13, 14]. These stressors also contribute to sharp declines in insect populations [15, 16] and rising outbreaks of pest species [17, 18]. As environmental pressures multiply and interact [19, 20], a central challenge in ecology is to understand and predict how ecosystems respond to change.

One important class of responses involves abrupt and difficult-to-predict changes that can trigger catastrophic regime shifts [4, 21–30]. These responses are often explained by models in which positive feedbacks generate alternative stable states and tipping points. Yet the extent to which such dynamics should be expected in real ecosystems remains unclear [31–34].

Ecosystem properties such as biomass or nutrient load can respond to gradual changes in environmental conditions (hereafter, stress) either by tracking the same trajectory during deterioration and recovery, or by following different paths. In the simplest case (FIG. 1a), deterioration and recovery follow the same smooth trajectory. In other cases (FIG. 1b), collapse and recovery occur at different stress levels because the system exhibits alternative stable states (hereafter, bistability). A collapse tipping point marks the stress level at which the system abruptly shifts to a degraded state, whereas a distinct recovery tipping point marks the lower stress level required to return to the original state. The separation between these paths is known as hysteresis and reflects strong positive feedbacks. Spatial heterogeneity can smooth the aggregate response by causing different patches to cross local thresholds at different stress levels [21, 35]. As a result, the system may appear to change gradually (FIG. 1c) even though collapse and recovery still follow different paths.

**FIG. 1.**
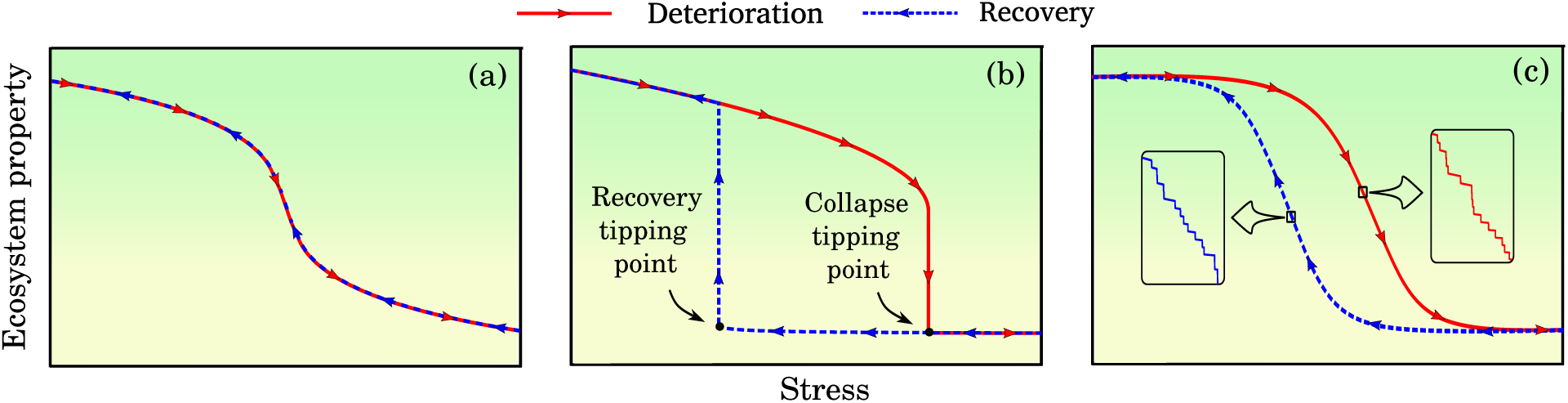
The potential responses of an ecological system to changes in environmental stress. Scenarios include: (a) a single, smooth trajectory during deterioration and recovery; (b) bistability, in which collapse and recovery follow different paths (hysteresis), producing abrupt shifts at distinct collapse and recovery tipping points; and (c) spatial heterogeneity, in which the aggregate response can appear smooth as different patches cross local thresholds at different stress levels, yet collapse and recovery still follow different paths. The seemingly smooth transition in (c) results from many small abrupt steps, as highlighted in the insets.

Ecological models of bistability, tipping points, and hysteresis, as illustrated in FIG. 1, have been applied to many systems, including lake eutrophication, insect outbreaks, desert vegetation, grazing dynamics, and species interactions in communities [23–30, 36, 37]. Over the past two decades, these models have received increasing attention, motivated by concerns that seemingly minor environmental changes could, in some settings, push ecosystems past a critical threshold and trigger an abrupt shift to a degraded state [21, 22, 38–40].

In management contexts, the analysis of these models has focused mainly on the sharp and difficult-to-reverse scenario: an ecosystem crosses a critical threshold, shifts abruptly to a degraded state, and then requires a much larger reduction in stress to recover. This possibility has strongly influenced management and conservation strategies, which often prioritize keeping systems away from putative tipping points [22, 31, 38, 41–48]. Examples include maintaining grazing pressure below critical levels to preserve vegetation cover [47, 49] and limiting nitrogen inputs to avoid ecosystem degradation [50, 51]. Major policy and conservation efforts have followed this paradigm, such as initiatives aimed at preventing desertification in the Sahel or mitigating harmful algal blooms in freshwater systems [11, 34, 52].

Although widely used in theory and management, the catastrophic shifts implied by these models have often eluded empirical detection [31, 33, 34, 53–58]. In parallel, theoretical concerns have emerged regarding the ability of classical catastrophic-shift models to capture real-world ecosystem dynamics [37, 59]. Many such models are well-mixed (spatially implicit) and deterministic, whereas natural ecosystems are shaped by complexities that can smooth or obscure abrupt transitions. These include spatial heterogeneity [21, 35], demographic stochasticity arising from random birth–death events in finite populations [37, 60], environmental stochasticity through temporally fluctuating conditions [61], and dispersal limitation that prevents complete spatial averaging and coupling. Yet despite growing recognition of these limitations of ecological theory, ecosystem management is still often framed around avoiding critical thresholds rather than anticipating gradual, cumulative, and spatially heterogeneous change [62, 63].

How likely are catastrophic shifts and alternative stable states given the ubiquity of stochasticity, spatial structure, and heterogeneity [64, 65]? More specifically, do tipping-point dynamics and bistability represent a generic risk wherever positive feedbacks exist, or do they persist only within a restricted region of parameter space?

Previous studies have highlighted several factors that can erode bistability and smooth transitions [21, 35–37, 59, 60, 66, 67]. However, what remains uncertain is how general this phenomenon is, how strongly it restricts the conditions for hysteresis, and through which dynamical routes hysteresis is lost. First, because most analyses focus on particular models or parameter choices, it is unclear whether similar effects arise across different ecological models, or whether the loss of hysteresis depends on specific model structures. Second, previous studies have not reconstructed full phase diagrams showing how collapse and recovery boundaries change across gradients of stress and complexity. Such phase diagrams are needed because they indicate not only whether tipping points occur, but also how wide the parameter region supporting bistability is, how stochasticity interacts with environmental stress, and whether complexity narrows the hysteretic region, or changes the qualitative nature of the transition. Third, stochasticity, spatial heterogeneity, and dispersal limitations act together in natural systems, yet their combined effects are rarely examined. If different forms of complexity compensate for one another, hysteresis may remain likely despite their individual effects; if their effects are additive, then even moderate levels of several complexities can jointly restrict the parameter space over which alternative stable states persist.

Here, we address these open questions by developing a systematic phase-diagram analysis of tipping-point dynamics and alternative stable states across four canonical ecological models. This approach allows us to map where hysteresis persists by tracking how collapse and recovery boundaries shift across gradients of environmental stress, stochasticity, spatial heterogeneity, and dispersal. It also allows us to test whether different sources of ecological complexity compensate for one another or accumulate. Across models, stochasticity and spatial heterogeneity restricted the parameter regions supporting hysteresis, while dispersal controlled their effective strength. When multiple complexity factors acted together, their effects were largely additive, suggesting that deterministic models may overstate the range of conditions under which tipping-point dynamics persist.

## RESULTS

To assess when alternative stable states and tipping points persist under realistic ecological complexity, we consider systems that can exhibit abrupt and irreversible transitions between a healthy state and a degraded state. We then map the phase diagram as a function of a stress parameter that pushes the system toward the degraded state and across the strength of stochasticity and spatial heterogeneity (FIG. 2).

**FIG. 2.**
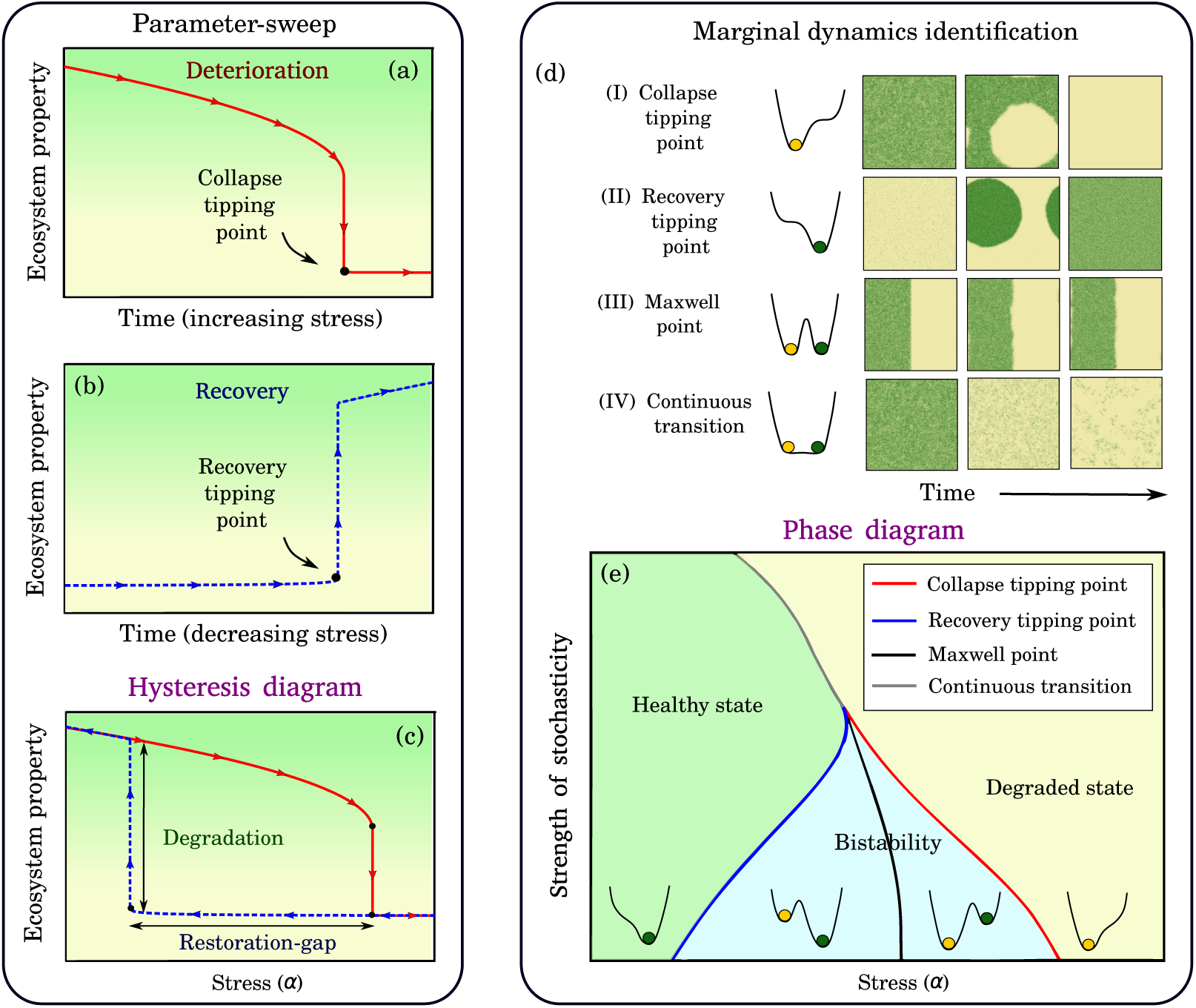
A framework for detecting tipping points and alternative stable states in stochastic, heterogeneous systems. (a) In the parameter-sweep approach, stress is increased gradually and the ecosystem property (e.g., biomass) is tracked; collapse is identified by an abrupt shift toward the degraded state. (b) In a downward sweep, stress is decreased gradually and recovery is identified by an abrupt shift toward the healthy state. (c) Combining upward and downward sweeps yields a hysteresis diagram, from which we quantify two indices: the *degradation index*, the maximal contrast in the ecosystem property between the two branches, and the *restoration-gap index*, the separation in stress between recovery and collapse. (d) In marginal dynamics identification, stress is held fixed and qualitative changes in stability and invasion dynamics are used to identify phase boundaries. Icons provide a stability-landscape schematic, used as a conceptual aid, in which valleys indicate locally stable states and ridges indicate unstable states. Examples for the desertification model include: (I) the *collapse tipping point*, where the healthy state loses stability and the mean time to collapse changes sharply; (II) the *recovery tipping point*, where the degraded state loses stability and establishment accelerates; (III) the *Maxwell point*, where invasion fronts stall because neither state has a systematic advantage; and (IV) a *continuous transition* under strong stochasticity, characterized by gradual decay to an absorbing state without hysteresis. (e) The resulting phase diagram summarizes qualitative regimes across stress levels and the strength of real-world complexity (e.g., stochasticity or heterogeneity). Transitions are defined in the legend. The corresponding representations of the system on a stability (“potential”) landscape [68], where valleys correspond to stable states and a tipping point occurs when a valley disappears, are shown in (d, e).

We use two complementary approaches to identify features in these phase diagrams. First, we perform slow upward and downward sweeps of the stress parameter, which yield hysteresis curves when collapse and recovery occur at different stress levels. These hysteresis curves provide two summary measures: a degradation index (DI), which quantifies the contrast between healthy and degraded states, and a restoration-gap index (RGI), which quantifies the irreversibility, i.e., the separation in stress between collapse and recovery. Second, we analyze system dynamics at fixed stress levels to locate qualitative landmarks in parameter space. This marginal-dynamics approach identifies the collapse and recovery tipping points and the Maxwell point, i.e., the stress level at which the two states are equally stable, in the sense that invasion fronts have no systematic tendency to advance in either direction.

We apply this framework to four canonical ecological models: a dryland desertification model, a lake pollution model, a grazing model, and an insect outbreak model. For brevity, we focus on the desertification and lake pollution models in the main text. Full descriptions of these models’ equations, parameters, and spatial and stochastic extensions are provided in the Methods. The remaining models are presented in the Supplementary Material.

Across all models, demographic and environmental stochasticity weaken hysteresis. As the strength of either source of stochasticity increases, hysteresis loops shrink and the abrupt jumps at collapse and recovery become smaller (FIG. 3a,b). Phase diagrams show how this erosion of hysteresis plays out across parameter space: regions in which only a healthy or only a degraded state exists expand, while the bistable region progressively contracts and can disappear under strong stochasticity (FIG. 3c,d).

**FIG. 3.**
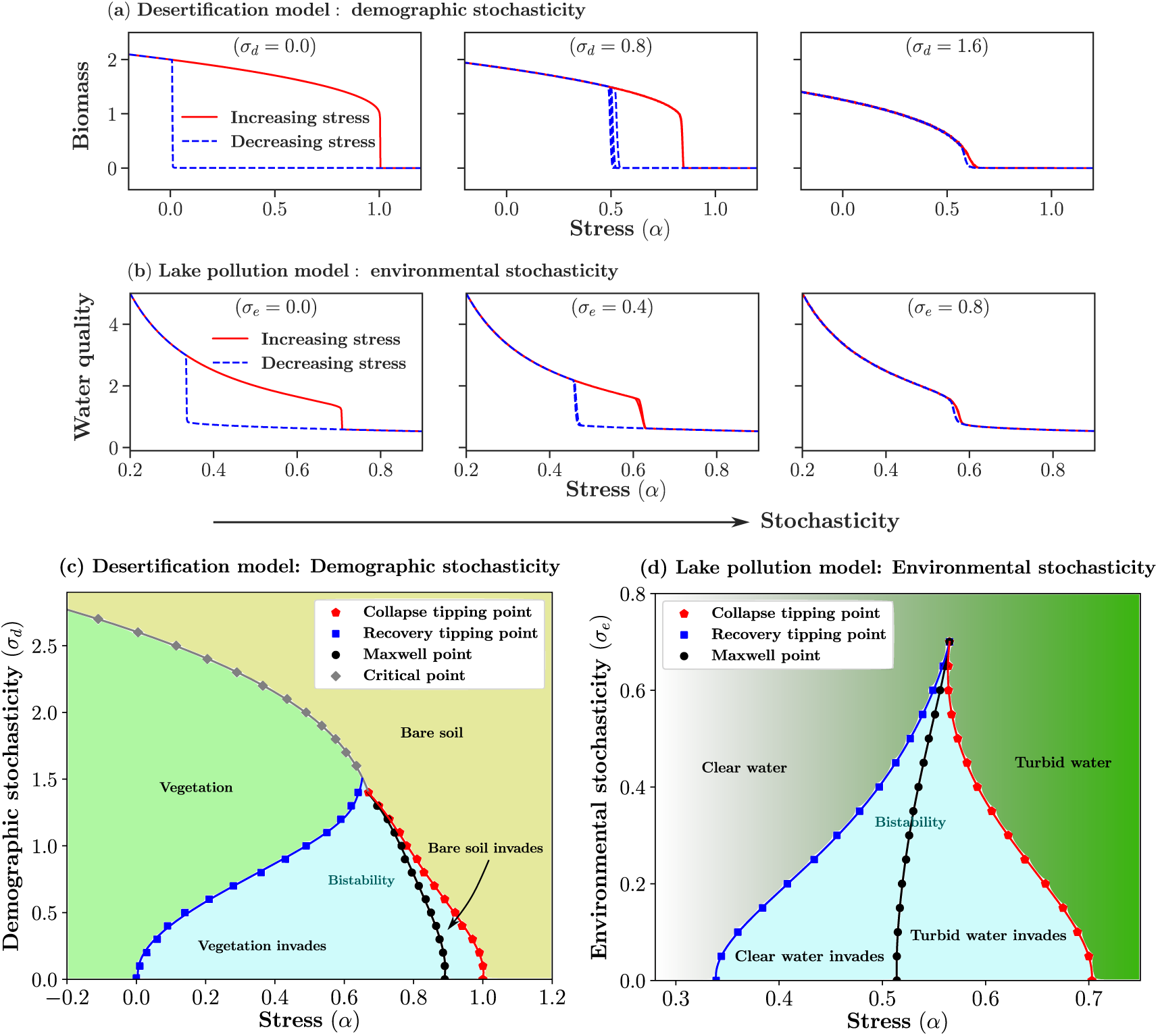
Demographic and environmental stochasticity reduce the prevalence of alternative stable states and abrupt transitions in the desertification and lake pollution models. (a) Hysteresis plots with increasing demographic stochasticity (*σ_d_*) in the desertification model. (b) Hysteresis plots with increasing environmental stochasticity (*σ_e_*) in the lake pollution model. The y-axes show the model state variable in arbitrary units: biomass in the desertification model and water quality (inverse algal biomass) in the lake pollution model. (c, d) Phase diagrams for (c) the desertification model and (d) the lake pollution model. Parameters: desertification *β* = 2, *γ* = 1, *D* = 1; lake pollution *β* = 1, *r* = 1, *δ* = 1, *D* = 1, *p* = 10. In all cases, *L*^2^ = 256^2^, and hysteresis plots show averages over 4 realizations.

An important distinction is apparent in FIG. 3c,d. In the desertification model, the zero-biomass degraded state is absorbing, i.e., a state from which the system cannot escape due to stochastic dynamics. Consequently, sufficiently strong stochasticity replaces the discontinuous, hysteretic shift with a continuous and reversible transition, in which biomass declines gradually to zero at the transition point [37]. In the lake pollution model, by contrast, neither alternative state is absorbing. As a result, under strong stochasticity the system exhibits a smooth crossover between phases, without a well-defined transition point. The phase diagrams further show that the bistable region does not shrink uniformly: in models with an absorbing low-biomass state, demographic stochasticity tends to expand the low-biomass region, whereas in models without an absorbing degraded state it behaves more similarly to environmental stochasticity (Supplementary Sections S4–S5; Figs. S42, S52).

The recovery boundary shows a counterintuitive effect. Although positive stress makes low biomass decay deterministically, demographic stochasticity can nucleate sufficiently large vegetated patches that grow and invade the degraded state. Thus, stochasticity can extend the active phase into a region where deterministic local dynamics predict biomass loss.

Under spatial heterogeneity, local habitats collapse or recover at different stress levels. This produces a third type of response: a transition that is gradual at the aggregate scale but still irreversible (FIG. 1c). Such transitions are not fully characterized by the degradation index or the restoration-gap index, because these measures quantify the size and irreversibility of hysteresis but not the sharpness of the aggregate response. We therefore introduce an abruptness parameter, defined as the maximum susceptibility along the hysteresis loop, max |*∂B/∂α*| (Supplementary O).

The abruptness depends on the interplay between heterogeneity and dispersal. Dispersal counteracts the effect of heterogeneity by coupling sites that would otherwise cross local thresholds independently, thereby reducing spatial fragmentation during the transition and increasing the abruptness of the regime shift. Panels (a) and (b) of FIG. 4 show that the abruptness is governed by Δ^2^*/D*, so when this parameter increases, the transition becomes gradual.

**FIG. 4.**
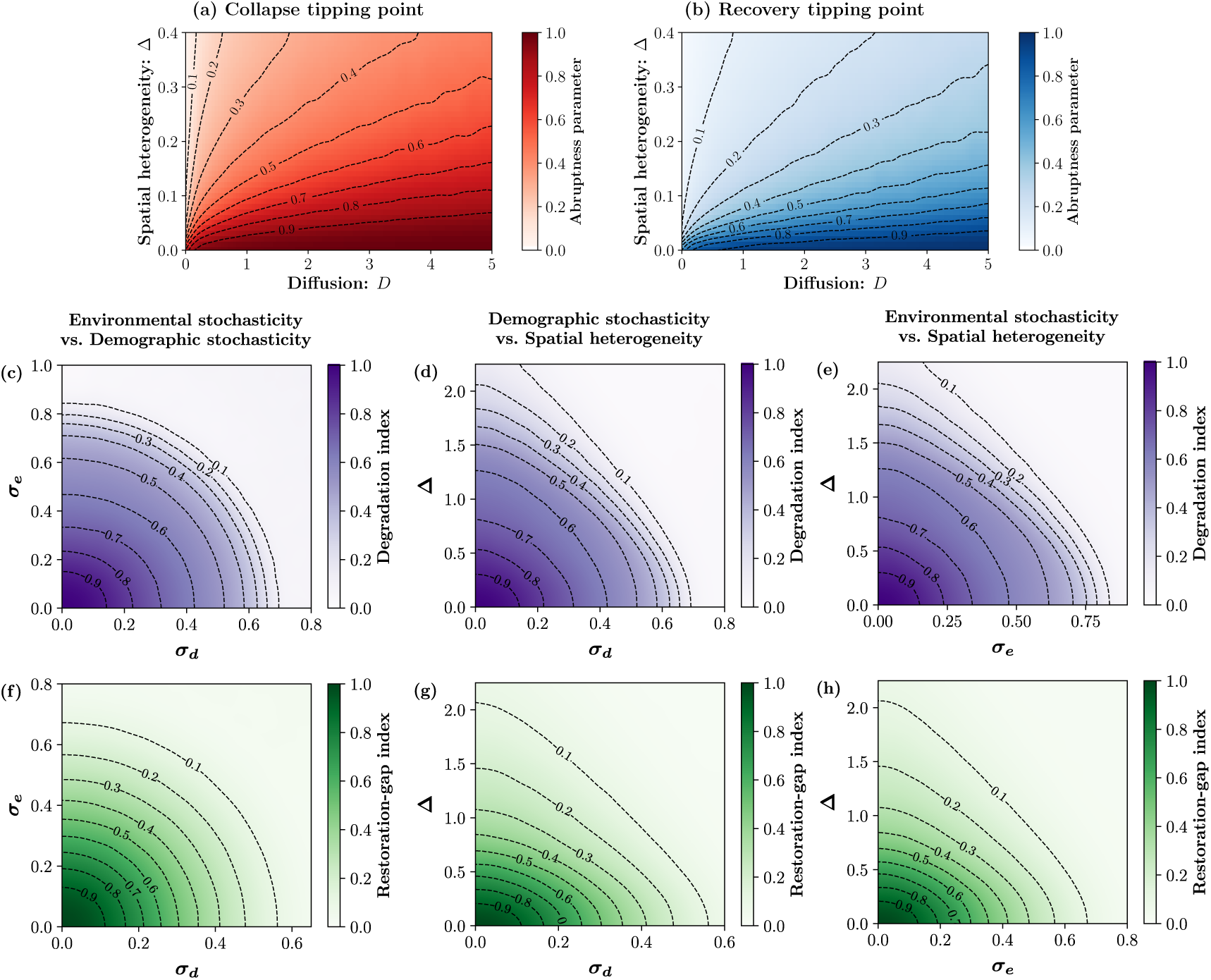
Interactions among real-world complexity factors and their consequences for hysteresis. First row (a,b): desertification model, showing abruptness parameter (normalized maximum of |*dB/dα*|) across spatial heterogeneity, Δ, and dispersal, *D*, for (a) the collapse tipping point and (b) the recovery tipping point. Second row (c–e): lake pollution model, showing the degradation index (contrast between healthy and degraded states). Third row (f–h): lake pollution model, showing the restoration-gap index (difference in stress, *α*, between the collapse and recovery tipping points). In both the second and third rows, panels (c, f), (d, g), (e, h) correspond to the parameter planes: (*σ_d_, σ_e_*), (*σ_d_,* Δ), and (*σ_e_,* Δ), respectively, where *σ_d_* is demographic stochasticity and *σ_e_* is environmental stochasticity. In all plots, darker shades indicate more abrupt transitions with stronger hysteresis, whereas lighter shades represent more gradual and more reversible responses. Black dashed contours mark equal index values (0.1–0.9). Across these parameter planes, the effects of different complexity factors are largely additive (for example, the indices in (c) often depend approximately on combinations such as 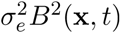), with substantial departures from additivity only for relatively large perturbations. Parameters: desertification *β* = 2, *γ* = 1, *L*^2^ = 128^2^, sweep rate *ɛ* = 4 × 10*^−^*^5^; lake pollution *β* = 1, *r* = 1, *δ* = 1, *D* = 1, *p* = 10, *L*^2^ = 256^2^, *ɛ* = 4 × 10*^−^*^5^.

Just as dispersal modifies the effective magnitude of spatial heterogeneity, it also modifies the effective magnitude of demographic and environmental stochasticity. With dispersal, local stochastic fluctuations are averaged across space, so the effective strength of stochasticity at the landscape scale diminishes as dispersal increases. We provide a comprehensive discussion of these relationships in the Supplementary Material. Broadly, we find that the effective strength of stochasticity is given by *σ*_eff_ = *σ/D^ζ^*, where *D* is the dispersal rate, *σ* denotes the strength of demographic or environmental stochasticity, and the exponent *ζ* typically lies between 1*/*3 and 1*/*2. Together, these scaling relationships provide a unified description of how dispersal rescales real-world complexities, enabling stochastic and heterogeneous effects to be compared across systems with different movement rates.

Notably, in natural ecosystems, stochasticity and spatial heterogeneity act simultaneously. Our analysis shows that their combined effects are largely additive (FIG. 4 c-h), leading to a joint reduction in both the degradation index and the restoration gap index.

These findings are general across both complexity factors and systems. The same qualitative erosion of hysteresis by stochasticity and spatial heterogeneity emerges in all models (desertification, grazing, outbreaks, lake pollution) despite their very different biological interpretations. Quantitative differences among models do exist, and results for each model are therefore reported in detail in the Supplementary Material.

## DISCUSSION

We have demonstrated that heterogeneity in time and space, together with demographic stochasticity, acts separately and collectively to restrict the conditions under which alternative stable states and tipping points persist. Across all four models, these real-world complexity factors consistently reduced the parameter ranges supporting abrupt collapse and difficult recovery, and their effects tended to reinforce rather than cancel one another. Thus, the presence of positive feedbacks in a deterministic model is not sufficient to infer that tipping-point dynamics will remain robust under realistic ecological complexity.

Despite this general pattern, the narrowing of the bistable region was not uniform. The direction in which phase boundaries shifted depended on model structure and on the type of stochasticity. In models with an absorbing bare-soil state, such as the desertification and grazing models, demographic stochasticity tends to favor bare soil because local fluctuations can drive patches toward extinction, where recovery is difficult. Yet the recovery boundary can shift in the opposite direction, because demographic fluctuations can also nucleate vegetated patches that expand into bare soil. In models without an absorbing degraded state, such as the lake pollution model, demographic stochasticity behaves more similarly to environmental stochasticity because no state can trap the system at zero abundance.

Notably, the largely additive nature of these effects means that different sources of complexity tend to accumulate rather than compensate for one another. Given the ubiquity of heterogeneity and stochasticity in natural ecosystems [65, 69], tipping points and alternative stable states should not be treated as automatic consequences of positive feedbacks. Rather, their persistence depends on whether the feedbacks generating bistability remain strong enough under the stochastic and heterogeneous conditions that characterize ecological systems.

### Toward more robust empirical tests of alternative stable states and tipping points

Empirical documentation of alternative stable states and tipping points remains ambiguous [31–34]. Comparatively strong evidence is often associated with relatively homogeneous systems such as shallow lakes [70], while some of the clearest demonstrations of collapse at a tipping point come from controlled experimental microcosms [71, 72]. If spatial and temporal heterogeneity erode bistability, then results from systems designed to minimize these factors cannot, on their own, establish that alternative stable states govern dynamics in more heterogeneous natural ecosystems. Even in one of the most widely discussed cases, forest versus savanna transitions [53, 73], recent studies suggest that environmental gradients, rather than bistability, may explain observed patterns [54, 74].

Part of this ambiguity reflects a broader gap between theoretical tipping-point models and empirical tests of their relevance. Much of the existing work has focused on identifying whether particular systems could exhibit alternative stable states or critical transitions, but these analyses often consider only part of the problem. Some studies calibrate deterministic parameters of process-based models [75], whereas others infer effective stability landscapes from empirical distributions [76]. In the latter case, the inferred potential is typically scaled by the unknown noise intensity, so deterministic forces and stochastic forcing are not separately estimated. Thus, even when empirical patterns are consistent with multiple attractors, it is often difficult to determine whether the underlying deterministic dynamics would support robust hysteresis once realistic fluctuations are included.

A complementary literature estimates demographic stochasticity, environmental variability [77], spatial heterogeneity [78], dispersal [79], or connectivity from ecological data. These quantities are central to real ecosystems, but they are usually estimated outside the framework of a calibrated nonlinear model with alternative stable states. As a result, the separate empirical advances needed to evaluate tipping-point robustness have rarely been integrated into a single analysis.

Our results indicate why this integration matters. We did not attempt to provide numerical parameter estimates for any particular ecosystem. Instead, we asked how the qualitative conclusions of canonical tipping-point models change when realistic sources of complexity are added. Within a unified theoretical framework, we showed that prior knowledge of stochasticity, environmental variability, spatial heterogeneity, and dispersal is not merely a technical refinement: these quantities can determine whether abrupt collapse and difficult recovery persist at all. The scaling relationships identified here provide a way to translate quantities that are often measured separately, such as dispersal and stochasticity, into effective parameters (e.g., *σ*_eff_ = *σ/D^ζ^*) that can be incorporated into analyses of catastrophic shifts. Together with the qualitative mechanisms described above, these relationships shift the empirical question from whether a deterministic model can generate alternative stable states to whether the resulting hysteresis remains robust under measured ecological complexity.

Answering this question requires several components that are rarely available together: a plausible deterministic skeleton for the focal system, estimates of demographic stochasticity and environmental variability, spatial heterogeneity, and information about dispersal or connectivity. To our knowledge, no empirical study has yet combined all these components in a way that allows the robustness of alternative stable states to be evaluated under stochasticity and spatial structure. Mesocosm experiments may provide one useful bridge, because they simplify the system while retaining stochasticity and spatial structure.

### Implications for interpreting ecosystem change and restoration

If stochasticity and heterogeneity can weaken hysteresis, then ecological patterns often interpreted as evidence for tipping points and alternative stable states may also warrant alternative explanations. Rapid ecological change does not necessarily imply that a system crossed a narrow tipping threshold; it may also reflect severe environmental change or repeated extreme events. Similarly, slow or incomplete recovery can emerge from slow life-history processes, dispersal limitation, species loss, or long transient dynamics, even without alternative stable states [80, 81].

This distinction is especially relevant for restoration and management, where the concept of alternative stable states strongly influences policy and practice [22, 24, 31, 42–44]. Land managers often use the term *state* as a practical label for distinct ecosystem conditions (e.g., grassland versus shrubland), an approach formalized in state-and-transition models [46]. This usage does not necessarily imply bistability in the theoretical sense of multiple equilibria. Recognizing this distinction does not reduce the practical value of state-and-transition models, but it clarifies that named management *states* should not automatically be interpreted as evidence for abrupt responses to gradual change or large restoration gaps caused by alternative stable states.

More broadly, managing ecosystems under global change may require moving beyond a purely threshold-based view, toward a perspective that recognizes gradual responses and appreciates the roles of stochasticity and spatial heterogeneity.

## ACKNOWLEDGMENTS

The authors acknowledge support from the Israel Ministry of Science (Grant No. 7034) and Israel Science Foundation (Grant 672/22) to N.D. N.M.S. thanks David Kessler for many helpful discussions. S.D. acknowledges the support of the Colman-Soref fellowship.

## METHODS

### Conceptual overview and models

We consider ecological systems that can exist in two contrasting states, a healthy state (such as a vegetated landscape or a clear lake) and a degraded state (such as bare soil or a turbid lake). The system is subject to environmental stress, represented by a control parameter, which may drive transitions between these states. Our objective is to determine under what conditions such systems exhibit alternative stable states (bistability) and hysteresis, and how the extent of these behaviors changes as real-world complexity increases.

We apply this approach to four canonical ecological models that have been widely used to study bistability and tipping points: a dryland desertification model [1, 2], a lake pollution model [3, 4], a grazing model [5], and an insect outbreak model [6]. For brevity, we focus on the first two in the main text and present the remaining models in the Supplementary Material.

In their simplest, well-mixed deterministic form, these models admit either a single stable state or bistability, depending on stress. In the desertification model, the well-mixed deterministic dynamics satisfy

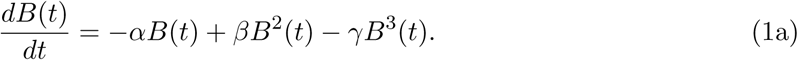

where *B*(*t*) denotes biomass density at time *t* and *α* represents environmental stress. Positive feedback is captured by the nonlinear growth term proportional to *β*, while *γ* limits growth at high biomass.

The lake pollution model is given by [3, 4]

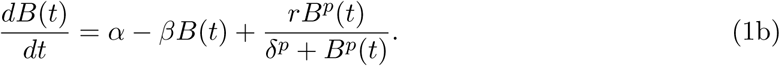

where *B*(*t*) represents nutrient density and *α* corresponds to external nutrient loading. The nonlinear recycling term introduces positive feedback and saturation, allowing bistability over an intermediate range of stress. Further details on these and the additional models are provided in the Supplementary Material (Section S1).

In spatial systems, alternative stable states need not be expressed as uniform equilibria across the entire domain. Different states can occupy different regions of the landscape, and their relative stability is reflected in their ability to expand or retreat through spatial invasion. Depending on stress level and dispersal, one state may invade the other. At a particular stress level, the invasion direction switches, separating conditions under which the healthy state expands from those under which the degraded state expands. This invasion balance point, often referred to as the Maxwell point, provides an additional boundary that complements collapse and recovery tipping points in spatial phase diagrams.

To incorporate real-world complexity, we extend the well-mixed models to spatially explicit systems and include four ubiquitous ecological ingredients. These ingredients are measurable in principle, and their joint quantification is exactly what is required to translate tipping-point theory into a falsifiable empirical assessment. Spatial heterogeneity captures persistent differences among local habitats that cause sites to experience different effective stress levels. Demographic stochasticity represents intrinsic variability arising from finite populations, whereas environmental stochasticity reflects temporal fluctuations in external drivers [7]. Lastly, dispersal couples neighboring locations, partially averaging local fluctuations and connecting heterogeneous patches, thereby modulating the effective influence of stochasticity and heterogeneity at the landscape scale.

To implement the above complexities within a minimal setting, we consider the dynamics of biomass on a two-dimensional square lattice of *L* × *L* sites. We label lattice sites by integer pairs (*m, n*) and denote the corresponding location by **x** ≡ (*m, n*) and the local biomass at a given time by *B*(**x***, t*). Demographic stochasticity is introduced, at each site, through an independent Wiener process with variance that scales like 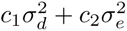, and the variance of the corresponding Wiener process for environmental stochasticity is 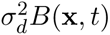. Spatial heterogeneity is implemented by modifying the local stress parameter as *α*(**x**) = *α* + Δ *ξ*(**x**), where Δ controls the strength of heterogeneity and *ξ*(**x**) represents fixed site-to-site variation, picked from a uniform distribution between [−1*/*2..1*/*2]. Spatial dispersal (diffusion) on the lattice is modeled as jumps to adjacent lattice sites at rate *D*.

Technically speaking, we integrate numerically a stochastic (Langevin) differential equation of the general form,

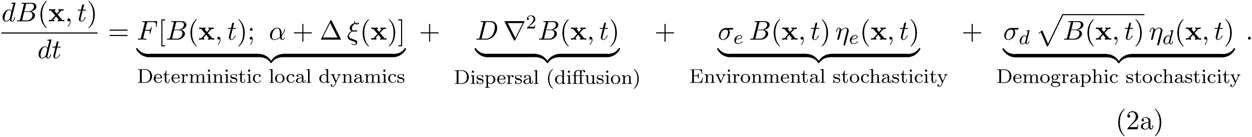

Here, *F* denotes the deterministic local dynamics of the corresponding well-mixed model (like Eqs.(1)), with stress replaced by the heterogeneous field *α* + Δ *ξ*(**x**). *η_e_* and *η_d_* are white noise processes, independent in space and time, and ∇^2^ denotes the discrete version of the Laplacian operator, i.e.,

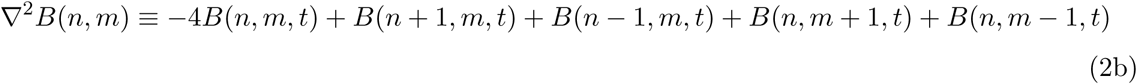

We simulate Eq.(2a) using an operator-splitting integration scheme for stochastic reaction–diffusion dynamics, following Refs. [8, 9]. At each time step, the local deterministic dynamics are updated together with environmental stochasticity and spatial heterogeneity, then the diffusive coupling among neighboring sites is applied by implementing the spectral method, and thereafter the demographic noise term is incorporated. The scheme preserves non-negativity, accurately captures absorbing states, and is widely used for spatial ecological models with intrinsic stochasticity. Additional simulation details are provided in the Supplementary Material (Section S2).

### Construction of phase diagram

To characterize tipping-point behavior across models and complexity levels, we map phase diagrams that summarize system dynamics across stress and the strength of real-world complexity (FIG. 2). Two complementary methods are implemented.

The first is a *parameter-sweep approach* (FIG. 2a–c), which is commonly used to visualize hysteresis by increasing or decreasing stress. A very slow sweep rate *ɛ* is implemented in these sweep protocols, to allow relaxation toward quasi-steady states. Tipping points then manifest themselves as discontinuous jumps in the biomass. Here, we extend this standard approach by introducing two quantitative indices: the *degradation index*, which measures the size of the jump, i.e., the contrast between the high- and low-state branches, and the *restoration-gap index*, which measures irreversibility through the separation in stress between collapse and recovery. Full sweep protocols and sensitivity analyses are reported in the Supplementary Material (Section S8).

The second method is *marginal dynamics identification* (FIG. 2d–e). By examining system behavior at a fixed stress level, this approach identifies the collapse and recovery tipping points, the Maxwell point separating opposing invasion directions, and, under strong stochasticity, a continuous transition in which hysteresis disappears.

This marginal dynamics method is implemented to the different phase boundaries as follows:

#### Collapse tipping point

The collapse tipping point *α_C_* is identified by initializing the system in the healthy state and monitoring the characteristic time required to transition to the degraded state over a suitable range of the stress parameter *α*. Below this tipping point, the characteristic time is very large; at and above the tipping point, it exhibits an abrupt change.

#### Recovery tipping point

The recovery tipping point *α_R_* is identified by initializing the system near the degraded state and quantifying establishment and growth of the healthy state. The recovery boundary corresponds to the stress level at which establishment becomes systematically favored, as determined from growth statistics averaged over stochastic realizations.

#### Maxwell point

The Maxwell point *α_MP_* is identified through invasion experiments in which adjacent healthy and degraded domains are initialized. The Maxwell point is defined as the stress level at which the invasion velocity vanishes, so that neither state systematically invades the other.

#### Continuous transition

In models with an absorbing degraded state, sufficiently strong demographic and environmental stochasticity can replace the discontinuous, hysteretic transition with a continuous one. We identify this boundary by the scale-free decay of healthy state toward the absorbing state.

For the system sizes and noise levels considered here, the predicted regime outcome is highly repeatable: independent stochastic realizations yield (nearly) identical hysteresis curves and phase boundaries, except very close to the transition lines where modest roughness is expected. In particular, we verified that sample-to-sample variability is small in our parameter ranges, and that the relevant indices converge with increasing system size and are largely insensitive to the sweep rates used. The full protocols and sensitivity analyses are reported in the Supplementary Material (Section S8).

Together, these two methods enable construction of full phase diagrams that delineate when bistability persists and how it is altered by real-world complexity. The use of two separate and independent methods also provides mutual cross-validation, since we require that both methods yield the same phase diagram, thereby increasing our confidence. See Supplementary S4 and S5 for details.

## Notes

### Competing Interest Statement

The authors have declared no competing interest.

