## Supplementary material for "Stochasticity and heterogeneity restrict tipping points and alternative stable states": supplementary_biorxiv.pdf

### Supplemental Information

#### S1: Details of Models

In this section, we describe the four models, the desertification, lake pollution, grazing, and insect outbreak models, in Subsections A, B, C, and D, respectively. We also provide remarks common to all models in Subsection E, including an overview of the potential-landscape framework for alternative stable states in Subsection F. Finally, we include a box (B1) that summarizes the key terminology used in the main text, together with a table (T1) listing the model-parameter values used throughout the paper.

**Box 1: Glossary:**

**Alternative stable states**: a situation in which an ecosystem can exist in distinct, stable regimes under the same environmental conditions (e.g., temperature or precipitation). This behavior is typically associated with positive feedback mechanisms, such as vegetation modifying local conditions in ways that promote further growth. When two such states coexist, the system is said to exhibit **bistability**. In analytical models, alternative stable states correspond to multiple stable equilibria separated by an unstable equilibrium.
*Synonyms*: `bistability`, `multistability`, `multiple equilibria`

**Hysteresis**: a phenomenon in which the response of a system to changes in an external driver depends on the direction of change, due to bistability. In ecological contexts, hysteresis implies that restoring a system to its former state requires reducing stress beyond the level at which collapse occurred.
*Synonyms*: `path dependence`, `irreversibility`

**Tipping point**: a threshold at which a small change in environmental conditions leads to an abrupt shift between alternative stable states. Tipping points occur at the boundaries of a bistable region, where one of the stable states loses stability. A **collapse tipping point** marks the loss of stability of the healthy state under increasing stress, whereas a **recovery tipping point** marks the loss of stability of the degraded state under decreasing stress.
*Synonyms*: `catastrophic shift`, `backward bifurcation`, `first-order transition`

**Maxwell Point**: a point at which two alternative stable states are equally favorable, so neither state invades the other. In spatial systems, it is characterized by vanishing invasion velocity between the two states.
*Synonyms*: `stall point`

**Absorbing State**: a state from which the system cannot escape under stochastic dynamics. In ecological models, this often corresponds to a zero-population state, such as bare soil in vegetation models, where demographic or environmental fluctuations have no effect. Stochastic systems tend to become trapped in absorbing states even when these states are unstable under deterministic dynamics.

**Continuous Transition**: a qualitative change in system behavior in which a stable and an unstable state merge as environmental conditions vary, leading to a smooth and reversible transition with no hysteresis. In spatial ecological systems, such transitions are typically characterized by a gradual decline of an order parameter to zero, rather than an abrupt jump. When the absorbing state is involved, continuous transitions often belong to the universality class of Directed Percolation [10, 11].
*Synonyms*: `second-order transition`, `transcritical bifurcation`

**Stability landscape (conceptual analogy)**: A qualitative representation in which the ecosystem state is depicted as a ball moving on a landscape shaped by environmental conditions. Valleys correspond to stable states and ridges to unstable states. A tipping point occurs when a valley disappears, forcing an abrupt transition to another state. This analogy is used as an interpretive aid without assuming the existence of a formal potential function.

##### A. Desertification model

Drylands occupy about 40% of Earth's land surface and are home to more than two billion people, roughly one-quarter of the global population [12, 13]. Understanding the mechanisms behind desertification in these regions is crucial, given their ecological and socio-economic importance. A simple yet insightful model describing the desertification process in arid and semi-arid ecosystems is given by Eq.(1a) in the main text.

In the stationary state, the model admits three solutions. A stable solution corresponds to the bare soil or the degraded state, given by $B_0 = 0$. Another stable solution represents the vegetated or healthy state, expressed as $B_s = \frac{1}{2\gamma}\left(\beta + \sqrt{\beta^2 - 4\alpha\gamma}\right)$, while the unstable solution is $B_u = \frac{1}{2\gamma}\left(\beta - \sqrt{\beta^2 - 4\alpha\gamma}\right)$. Here, $\alpha$, $\beta$, and $\gamma$ are model parameters that control biomass dynamics (also act as a stress parameter), positive feedback strength, and resource limitation, respectively.

For this model, one can compute the tipping points and the Maxwell point exactly in terms of the model parameters. The collapse tipping point, where vegetation collapses to bare soil, occurs at $\alpha_C = \beta^2/4\gamma$, which follows directly from the condition $\sqrt{\beta^2 - 4\alpha\gamma} \geq 0$.

The recovery tipping point, where vegetation can spontaneously reestablish, occurs at $\alpha_R = 0$, and is obtained by analyzing the stability of the $B_0 = 0$ solution. Specifically, taking the derivative of the right-hand side of the deterministic model with respect to $B$, and then evaluating it at $B = 0$, yields $-\alpha$. Clearly, as $\alpha$ changes from positive to negative, this quantity changes sign, indicating that the solution loses stability.

The Maxwell point, where the vegetated and bare soil states are equally favorable, is located at $\alpha_{MP} = 2\beta^2/9\gamma$ (the derivation will be discussed in Subsection F). To ensure the existence of alternative stable states (i.e., bistability), the positive feedback coefficient must satisfy $\beta > 0$.

##### B. Lake pollution model

A recent study predicts that, by 2050, half of the world's lakes will be eutrophic and another 5% hypereutrophic, leaving only 21% oligotrophic and 24% mesotrophic [14]. Clearly, these ecosystems face growing threats from pollution and nutrient loading [15]. A simple but insightful model capturing the pollution dynamics of a lake is presented in Eq.(1b) in the main text.

As described in the main text, the model incorporates the parameters $\alpha$, $\beta$, $r$, $\delta$, and $p$. The parameter $\alpha$ represents the external pollution loading rate (acting as a stress parameter), $\beta$ controls the natural recovery rate of the lake, $r$ measures the recycling strength of pollutants within the lake, and $\delta$ denotes the nutrient density at which the recycling rate reaches $r/2$. The exponent $p \geq 2$ governs the steepness of the recycling process and can vary from 2 for deep cold lakes to 20 for shallow warm lakes [16].

In the stationary state, the model admits three possible solutions. For small values of $\alpha$, the system settles into an oligotrophic equilibrium, characterized by clear and healthy water. For large values of $\alpha$, it shifts to a eutrophic equilibrium, marked by turbid and degraded water. Separating these two stable states is an unstable intermediate equilibrium, which defines the boundary between their respective basins of attraction.

In contrast to the desertification model, there is no analytical expression for the tipping and Maxwell points, as the deterministic model in the stationary state yields a polynomial equation of degree $p+1$. In this case, for a fixed set of model parameters, we plot $\alpha + \frac{rB^p}{\delta^p + B^p}$ and $\beta B$ versus $B$. The intersection of these two curves yields either one or three solutions. One intersection corresponds to a single stable solution (representing either an oligotrophic or eutrophic equilibrium), while three intersections indicate two stable solutions separated by an unstable equilibrium. By varying the stress parameter $\alpha$, we find the specific points at which the number of intersections switches between one and three, thus identifying the tipping points of the model. To calculate the Maxwell point numerically, we measure the difference in the depths of the potential landscape associated with the two stable solutions (see Subsection F for further details on the potential landscape framework).

##### C. Grazing and Overgrazing in Rangelands

Managed grazing spans more than 25% of the global land area, covering a larger geographical extent than any other land use [17]. The dynamics of biomass in a grazing system is commonly described by Noy-Meir's model [5]. In this model, the biomass $B(t)$ of the vegetation at time $t$ follows the equation:

$$\frac{dB(t)}{dt} = rB\left(1 - \frac{B}{K}\right) - \frac{\alpha B}{\delta + B}. \tag{S11}$$

The growth of biomass follows a logistic function where $r$ is the maximal growth rate, and $K$ is the carrying capacity (the equilibrium biomass without grazing). The consumption rate of the grazer is a Michaelis-Manten function of biomass, with $\alpha$ as the maximal consumption rate of the domestic grazers (acting as a stress parameter) and $\delta$ being the half-saturation constant (the biomass level where consumption is half of its maximal rate).

In the stationary state, the above equation has three solutions. One stable solution is the bare soil state, $B_0 = 0$. Another stable solution corresponds to vegetation, $B_s = \frac{K-\delta}{2} + \frac{1}{2}\sqrt{(K+\delta)^2 - \frac{4\alpha K}{r}}$. There is also one unstable solution, $B_u = \frac{K-\delta}{2} - \frac{1}{2}\sqrt{(K+\delta)^2 - \frac{4\alpha K}{r}}$. The collapse tipping point can be computed as $\alpha_C = \frac{r}{4K}(K+\delta)^2$, which follows directly from the condition $\sqrt{(K+\delta)^2 - \frac{4\alpha K}{r}} \geq 0$. The recovery tipping point is given by $\alpha_R = \delta r$, which is obtained by analyzing the stability of the $B_0 = 0$ solution. The Maxwell point ($\alpha_{MP}$) is numerically obtained by plugging the stable solution of vegetation into this equation (see in Subsection F):

$$-\alpha\delta \log\left(\frac{\delta}{\delta + B}\right) = \alpha B + \frac{rB^3}{3K} - \frac{rB^2}{2}$$

To obtain alternative stable states, we require $K > \delta$ in this case.

##### D. Insect Outbreaks model

Insect outbreaks, particularly by forest pests such as bark beetles and defoliators, impact millions of hectares of forests globally each year, leading to significant ecological and economic consequences [18]. The most widely used model for insect outbreaks in forests was developed by Ludwig, Jones, and Holling [6, 19]. Although it describes a different biological phenomenon, the structure of this model closely resembles that of the grazing model. In both cases, the population biomass $B$ grows logistically and is subject to consumption, with the consumer in the outbreak model playing a role analogous to the grazer in the grazing model. The governing equation for this system is

$$\frac{dB(t)}{dt} = rB\left(1 - \frac{B}{K}\right) - \frac{\alpha B^2}{\delta^2 + B^2}. \tag{S12}$$

However, due to a small but crucial difference in the consumption term—being sigmoidal in this model—there is no stable steady state at zero biomass. Instead, similar to the lake pollution model, the system exhibits two alternative stable states: a high-density outbreak state and a low-density refuge state. Additionally, there are two unstable equilibria: one at $B_0 = 0$ and another between the two stable states. Excluding the trivial solution at $B_0$, the non-trivial stationary states can be determined by numerically solving

$$B\left(\delta^2 + \frac{\alpha K}{r}\right) - KB^2 + B^3 = K\delta^2.$$

Unlike the grazing model, the tipping and Maxwell points in this case are computed numerically, following a procedure similar to that used in the lake pollution model.

##### E. General Remarks common for all models

Apart from the model-specific details, we can make the following general comments:

To develop a simple model of alternative stable states that exhibits a catastrophic shift, three essential components are required. The first is the growth and/or mortality rates ($\alpha$ in desertification and lake pollution models and $r$ in grazing and insect outbreak models), which govern the increase or decline of population/biomass over time in the absence of additional complexities. The second is a finite carrying capacity ($K$ and $\gamma$) that limits indefinite biomass growth by imposing an upper bound within a finite-sized system. The third and the most crucial element is a positive feedback mechanism ($\beta$), which introduces memory into the system's dynamics, eventually giving rise to hysteresis and the emergence of alternative stable states in the stationary regime.

It is clear from the description of the models that the desertification and grazing models are analogous in the sense that both possess an absorbing state ($B_0 = 0$). In contrast, the insect outbreak model closely resembles the lake pollution model, as both exhibit two non-zero stable stationary solutions. In this paper, we examine how these similarities and differences among the models influence the results at both qualitative and quantitative levels.

##### F. Potential landscape framework for understanding alternative stable states

The concept of alternative stable states or bistability in each model can be intuitively visualized using a potential landscape framework [20]. In this framework, the system state is represented by a ball rolling in a landscape defined by a potential function. The minima of this potential landscape correspond to stable states, and the position in which the ball eventually settles depends on its initial location—i.e., the initial condition of the system.

The potential landscape $V(B)$ can be derived by integrating the negative of the right-hand side of the deterministic equation with respect to the biomass variable $B$ [20]. For instance, in the desertification model, the effective potential takes the form:

$$V(B) = \frac{\alpha B^2}{2} - \frac{\beta B^3}{3} + \frac{\gamma B^4}{4}. \tag{S13}$$

As the stress parameter $\alpha$ is varied while keeping $\beta$ and $\gamma$ fixed, the shape of the potential landscape undergoes qualitative changes. For $\alpha > \alpha_C$, the landscape exhibits a single minimum at $B = 0$, representing a degraded, bare-soil state with no vegetation. In this regime, the healthy vegetated state (i.e., the non-zero stationary solution of the model) is no longer stable, as the potential minimum corresponding to it has vanished. Conversely, for $\alpha < \alpha_R$, the landscape also becomes monostable, but now with a single minimum corresponding to the healthy vegetated state. The degraded state is unstable in this regime, and the system invariably recovers vegetation regardless of its initial condition. Between these two tipping points, i.e., for $\alpha_R \leq \alpha \leq \alpha_C$, the potential landscape features two minima separated by a maximum (unstable equilibrium). This bistable regime supports both the healthy and degraded states as locally stable solutions, with the system's long-term behavior determined by its initial condition.

Within this bistable range, there exists a specific value of the stress parameter, known as the Maxwell point, denoted by $\alpha_{MP}$, at which the depths of the two potential minima become equal, i.e., $V(B_0) = V(B_s)$. For the desertification model, substituting $B_0$ and $B_s$ into the expression for $V(B)$ yields an analytical expression for the Maxwell point: $\alpha_{MP} = \frac{2\beta^2}{9\gamma}$. In the grazing model, an equation based on $V(B_0) = V(B_s)$ can be derived and solved numerically to obtain the Maxwell point. For the other two models, we compute the Maxwell point numerically by evaluating the difference in potential depth between the two stable states; the value of the stress parameter at which this difference changes sign is identified as the Maxwell point.

For $\alpha_R < \alpha < \alpha_{MP}$, although both states are stable, the bare-soil state corresponds to a shallower potential well and is thus metastable relative to the vegetated state in desertification model. Conversely, for $\alpha_{MP} < \alpha < \alpha_C$, the vegetated state becomes metastable, as its potential well is shallower than that of the bare-soil state.

In spatially explicit models, that is, when the system is placed on a lattice with dispersal (diffusion), these two ranges of the stress parameter have important implications. In such systems, one stable state can invade the other within these regimes. Specifically, for $\alpha_R < \alpha < \alpha_{MP}$, vegetation invades the bare-soil state, whereas for $\alpha_{MP} < \alpha < \alpha_C$, the bare-soil state invades the vegetation in case of desertification model.

| **Model** | **Model Parameters** |
|---|---|
| Desertification | $\beta = 2,\ \gamma = 1$ |
| Lake pollution | $\beta = r = \delta = 1,\ p = 10$ |
| Grazing & Insect outbreak | $r = 1,\ K = 10,\ \delta = 0.5$ |

TABLE T1. Summarized parameters for all models used throughout the paper.

#### S2: Simulating the dynamics: operator-splitting method

To numerically solve the nonlinear stochastic partial differential equation [Eq. (2a) in the main text], we employ an operator-splitting scheme following Weissmann *et al.* [8], in which each contribution is integrated separately in a sequential manner. This strategy enables efficient and stable treatment of the different deterministic and stochastic components. At each time step, the system is advanced by performing the following four steps:

**Step I**: We begin by integrating the deterministic component of the equation. At each lattice site, the biomass is updated using the explicit Euler method over an infinitesimal time increment

$\Delta t$:

$$B_1(\mathbf{x}, t+\Delta t) = B_0(\mathbf{x}, t) + \Delta t \left[\text{Deterministic components with } \alpha(\mathbf{x}) \to \alpha + \Delta\xi(\mathbf{x})\right]. \tag{S21}$$

During this step, spatial heterogeneity is incorporated by modifying the local stress parameter as $\alpha(\mathbf{x}) = \alpha + \Delta\xi(\mathbf{x})$, where $\xi(\mathbf{x})$ is independently drawn from a uniform distribution over $[-1/2, 1/2]$ for each site.

**Step II**: Using the result of Eq.(S21) as input, we next incorporate environmental stochasticity by applying the analytical solution to a geometric Brownian motion of the form $dB_t = \sigma_e B\, dW_t$, where $dW_t$ is a Wiener process with zero mean and variance $t$. Over a time interval $\Delta t$, the biomass at each site evolves according to (Itô integration):

$$B_2(\mathbf{x}, t+\Delta t) = B_1(\mathbf{x}, t+\Delta t) \cdot \exp\left[\sigma_e \Delta W - \frac{1}{2}\sigma_e^2 \Delta t\right], \tag{S22}$$

where $\Delta W \sim \mathcal{N}(0, \Delta t)$ is a normally distributed random variable of zero mean and variance $\Delta t$. To maintain biological realism, we enforce non-negativity by resetting any negative biomass values after Steps I and II to zero.

**Step III**: We then address spatial dispersal using a spectral method. First, the Fourier transform of $B_2(\mathbf{x}, t+\Delta t)$ is computed:

$$B_2(\mathbf{k}, t+\Delta t) = \mathcal{F}\left[B_2(\mathbf{x}, t+\Delta t)\right],$$

where $\mathbf{k} = (k_x, k_y)$ is the wavevector in Fourier space. The effect of diffusion over time $\Delta t$ is then applied multiplicatively in spectral space:

$$B_3(\mathbf{k}, t+\Delta t) = B_2(\mathbf{k}, t+\Delta t) \cdot \exp\left[2D\Delta t(\cos k_x + \cos k_y - 2)\right],$$

followed by an inverse Fourier transform to return to real space:

$$B_3(\mathbf{x}, t+\Delta t) = \mathcal{F}^{-1}\left[B_3(\mathbf{k}, t+\Delta t)\right]. \tag{S23}$$

**Step IV**: Finally, demographic stochasticity is implemented using the exact solution of the corresponding Fokker–Planck equation [21], with random deviates generated via the procedure of Dornic *et al.* [9]. For each site, a Poisson-distributed random variable is first drawn with mean $\lambda B_3(\mathbf{x}, t+\Delta t)$, where $\lambda = 2/(\sigma_d^2 \Delta t)$. The resulting value is then used as the shape parameter of a Gamma distribution with unit scale. A Gamma variate is sampled and subsequently rescaled by $1/\lambda$:

$$B_4(\mathbf{x}, t+\Delta t) = \frac{1}{\lambda}\,\text{Gamma}\left(\text{Poisson}\left[\lambda B_3(\mathbf{x}, t+\Delta t)\right], 1\right). \tag{S24}$$

This algorithmic step captures intrinsic noise in finite populations and ensures that the variance in biomass scales linearly with the current biomass, consistent with the underlying demographic stochasticity.

We repeat Steps I through IV iteratively to advance the biomass dynamics over time using a fixed time step of $\Delta t$ throughout the simulations. In this paper we use $\Delta t = 0.05$, and 0.1. These time steps are chosen to balance numerical stability with computational efficiency while preserving the fidelity of the stochastic and spatial dynamics.

#### S3: The degradation index (DI) and the restoration gap index (RGI)

In this section we describe the methodology used to compute the degradation index (DI) and the restoration–gap index (RGI). For each model and parameter set, we first construct the full hysteresis diagram using the parameter sweep method. Once the hysteresis diagram is constructed, DI and RGI are computed as follows:

**Degradation index:** The DI is defined as the largest biomass difference between the deterioration and recovery branches, evaluated at the same stress level $\alpha$ (violet dashed line in FIG. S31). Operationally, we construct a dense, uniform grid in $\alpha$ over the overlap of the two branches. For each grid point, we evaluate $B$ on both branches, compute the absolute vertical separation, and take the maximum over the grid as the DI.

**Restoration–gap index:** The RGI is defined as the largest stress difference between the deterioration and recovery branches, evaluated at the same biomass $B$ (green dashed segment in FIG. S31). Again, we construct a dense, uniform grid in $B$ over the range common to both branches. For each grid point, we estimate $\alpha$ on both branches, compute the absolute horizontal separation, and take the maximum over the grid as the RGI.

#### S4: Role of Demographic stochasticity

In this section, we examine in detail how demographic stochasticity affects alternative stable states and the associated collapse and recovery tipping points. To focus exclusively on these effects, we include the demographic noise and spatial dispersal alongside the deterministic components of the

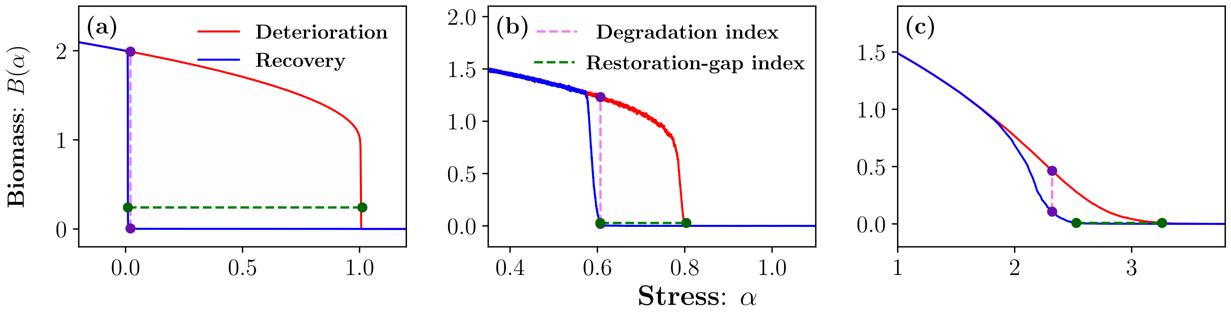


FIG. S31. Biomass $B(\alpha)$ as a function of stress $\alpha$, obtained by a parameter sweep at rate $\epsilon = 10^{-5}$ for the desertification model: (a) $\sigma_d = 0.0$, $\Delta = 0.0$; (b) $\sigma_d = 1.0$, $\Delta = 0.0$; (c) $\sigma_d = 0.0$, $\Delta = 9.0$; other parameters fixed at $\beta = 2$ and $\gamma = D = 1$ with $L^2 = 256^2$. The deterioration (increasing $\alpha$) and recovery (decreasing $\alpha$) branches are shown in red and blue, respectively. The degradation index is indicated by the violet dashed *vertical* line, and the restoration–gap index by the green dashed *horizontal* line.

dynamics, i.e.,

$$\frac{dB(\mathbf{x},t)}{dt} = \text{Deterministic components} + \underbrace{\sigma_d \sqrt{B(\mathbf{x},t)}\,\eta_d(\mathbf{x},t)}_{\text{Demographic stochasticity}} + \underbrace{D\ \nabla^2 B(\mathbf{x},t)}_{\text{Spatial dispersal}}\ .$$

In what follows, we illustrate how demographic stochasticity, quantified by its strength $\sigma_d$, modifies the hysteresis diagrams constructed using the parameter-sweep method (Subsection G). We then present phase diagrams in the $(\alpha, \sigma_d)$ plane obtained using the marginal dynamics identification method (Subsection H), highlighting how demographic stochasticity alters the locations of different transition points. In the same subsection (H), we briefly discuss the similarities and differences among phase diagrams obtained from different models at a qualitative level. We also highlight the consistency between the parameter-sweep and marginal dynamics identification methods. Finally, in Subsection I, we discuss how diffusion modulates demographic stochasticity and compute an effective stochasticity $\sigma_{d,\text{eff}}$ combining both $\sigma_d$ and $D$.

#### G. Hysteresis Diagrams: Parameter sweep method

In this subsection, we plot hysteresis diagrams for the four models with increasing demographic stochasticity ($\sigma_d$) to show that increasing the strength of $\sigma_d$ progressively softens an abrupt, effectively irreversible transition into a smooth, reversible one (see Fig. S41).

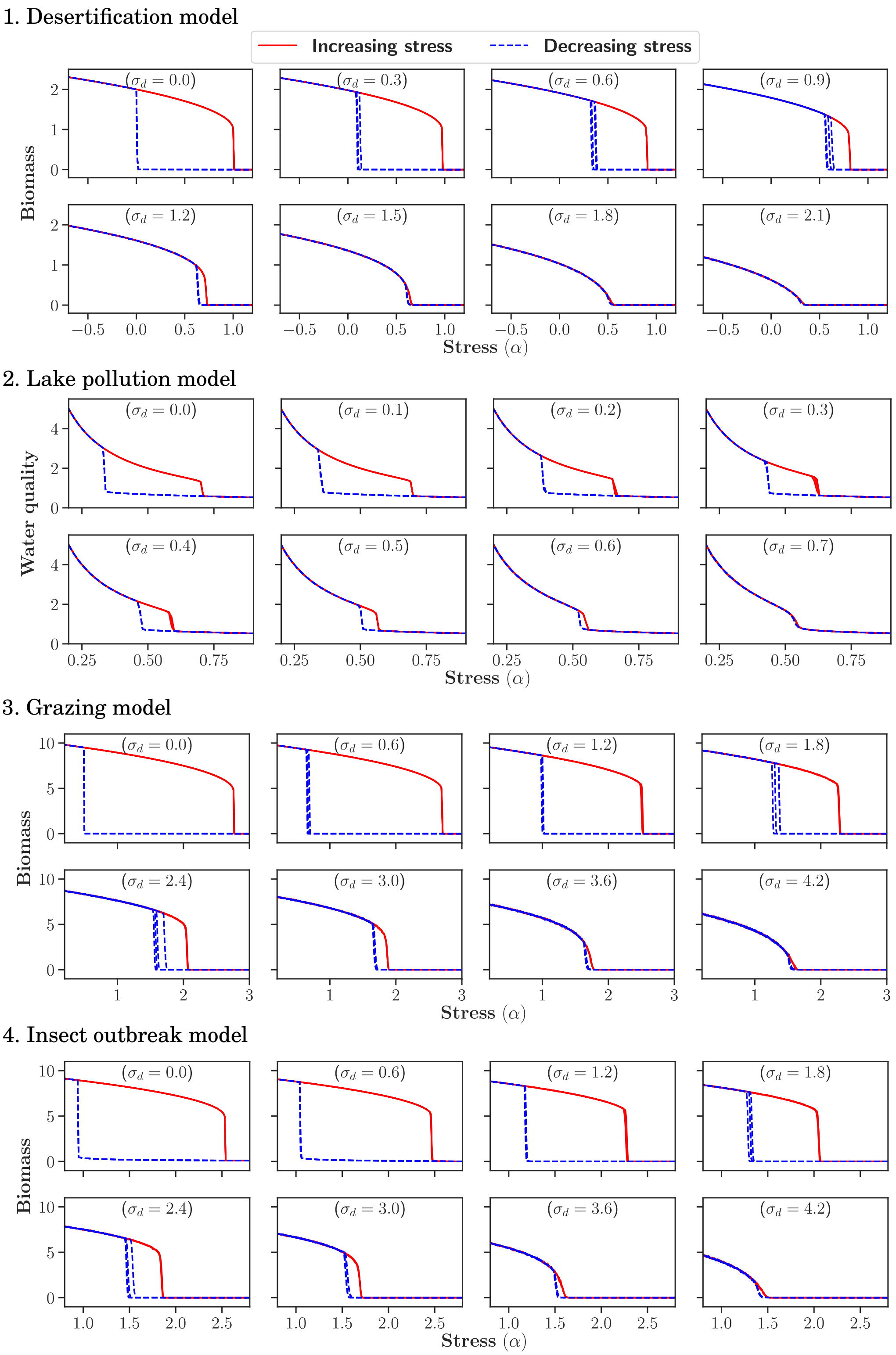


FIG. S41. Hysteresis plots for the 1. desertification, 2. lake pollution, 3. grazing, and 4. insect outbreak models. For each model, we show eight diagrams corresponding to increasing levels of demographic stochasticity, with four realizations for each diagram. Model parameters are listed in Table T1; additionally, we use dispersal rate $D = 1$, sweep rate $\epsilon = 10^{-5}$, and system size $L \times L = 256^2$.

##### H. Phase Diagrams: Marginal dynamics identification

In this subsection, we present complete phase diagrams in the $(\alpha - \sigma_d)$ plane obtained through marginal dynamics identification for all four models (see FIG. S42). In all cases we identify the Maxwell (stall) point and the two tipping points, and see how these points merge as stochasticity increases in strength.

In the strong stochasticity phase, one would like to distinguish between two types of dynamics. In the cases of desertification and grazing (FIG. S42(a) and (c)), one of the alternative steady state, the bare soil state, is absorbing, i.e., it is unaffected by stochasticity. Therefore, as stress increases, the steady state density decreases until it hits zero. This is a continuous phase transition (marked as a "critical point" in the relevant panels) with critical exponents that belong to the directed percolation class [2]. On the other hand, the lake pollution dynamics (FIG. S42(b)) does not have an absorbing state, as both stationary state solutions are non-zero; consequently, no critical point appears in its strong-stochasticity part of its phase diagram.

Interestingly, the insect outbreak model exhibits a critical point under high levels of demographic stochasticity, even though it does not exhibit an absorbing stationary state in the deterministic limit. This behavior arises because demographic stochasticity effectively transforms the lowest non-zero stationary state into an absorbing state beyond a critical threshold. The corresponding threshold is indicated by the dark cyan line in FIG. S42(d): below this line, the phase diagram resembles that of the lake pollution model, whereas above it, the system's behavior aligns with that of the desertification and grazing models. To identify this line, we consider an initially empty system containing a single degraded patch and determine the critical value of $\sigma_d$ at which the degraded state becomes extinct for a fixed set of stress and other parameters.

In all diagrams of FIG. S42, we have also included tipping points obtained using the parameter-sweep method (dotted and dashed lines in each diagram). For each model, the tipping points are shown for two different sweep rates, $\epsilon = 10^{-5}$ and $10^{-4}$, demonstrating that the two approaches complement each other: as $\epsilon$ decreases, the tipping points obtained from the parameter-sweep method converge toward those derived from marginal dynamics identification.

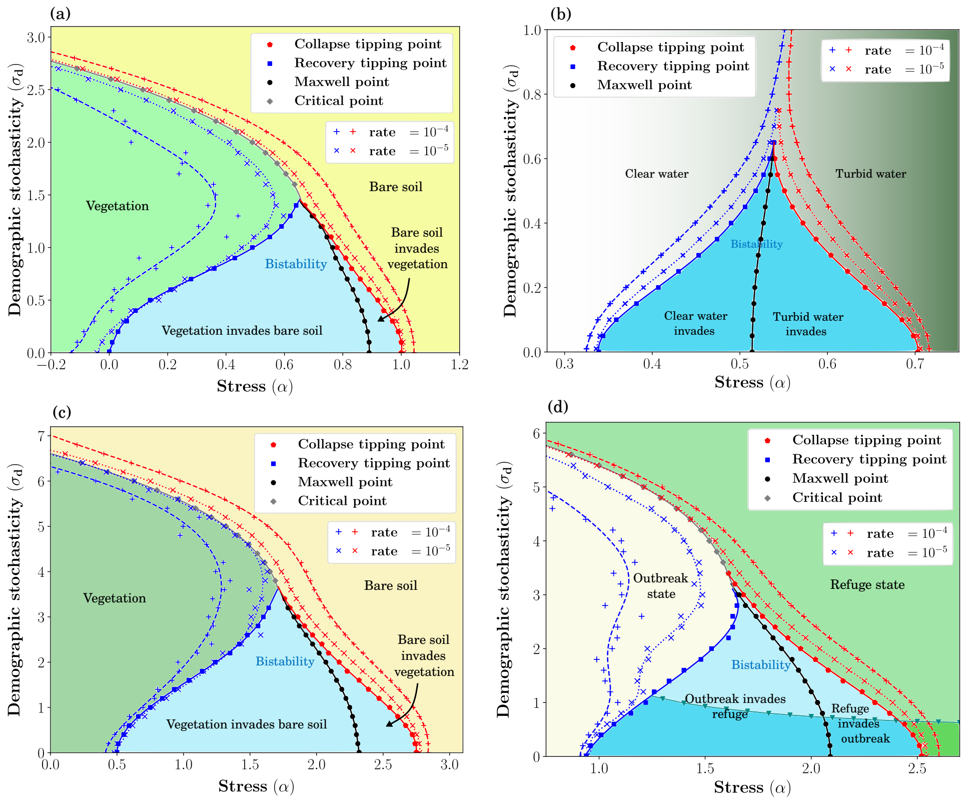


FIG. S42. **Phase diagrams in the $(\alpha - \sigma_d)$ plane for (a) desertification, (b) lake pollution, (c) grazing, and (d) insect outbreak models.** Solid lines denote phase boundaries obtained using the marginal dynamics identification method. The regions shaded in light cyan correspond to bistable domains where one of the stationary states is absorbing, while the dark cyan regions indicate bistability between two non-zero stationary states. Dotted $(\times)$ and dashed $(+)$ lines show tipping-point trajectories obtained from the parameter-sweep method for rates $\epsilon = 10^{-5}$ and $\epsilon = 10^{-4}$, respectively. Model parameters are listed in Table T1 with additional dispersal rate $D = 1$ and system size $L \times L = 256^2$.

#### I. Effect of diffusion on demographic stochasticity

As noted in the main text, increasing stochasticity, for example, demographic stochasticity with strength $\sigma_d$ enhances the spatial roughness of biomass, whereas stronger dispersal (characterized by diffusion constant $D$) counteracts this by redistributing biomass and smoothing spatial patterns. These two processes therefore act in opposition, suggesting the existence of an effective stochasticity $\sigma_{d,\text{eff}}$ that combines both $\sigma_d$ and $D$. Based on numerical simulations, we have found $\sigma_{d,\text{eff}} = \sigma_d / D^{\zeta}$.

To examine this hypothesis numerically, we check for a data collapse, i.e., we expect the phase diagrams plotted for different values of $D$ and $\sigma_d$ to collapse once $\zeta$ is chosen appropriately. These collapses are demonstrated in FIG.S43.

For models with two non-zero stationary solutions at the deterministic limit (e.g., the lake–pollution model), setting $\zeta \approx 1/2$ yields a clear data collapse for $D = 1/2, 1, 2$ within the bistable regime (see FIG.S43(b)). For models with an absorbing stationary solution at the deterministic limit (e.g., the desertification and grazing models), the collapse is well described by $\zeta \approx 1/3$ within the bistable regime (see FIG.S43(a),(c)). As explained, in the insect–outbreak model, demographic stochasticity can effectively transform the lower nonzero stationary state into an absorbing state; accordingly, the data collapse follows $\zeta \approx 1/3$ rather than 1/2 (see FIG.S43(d)).

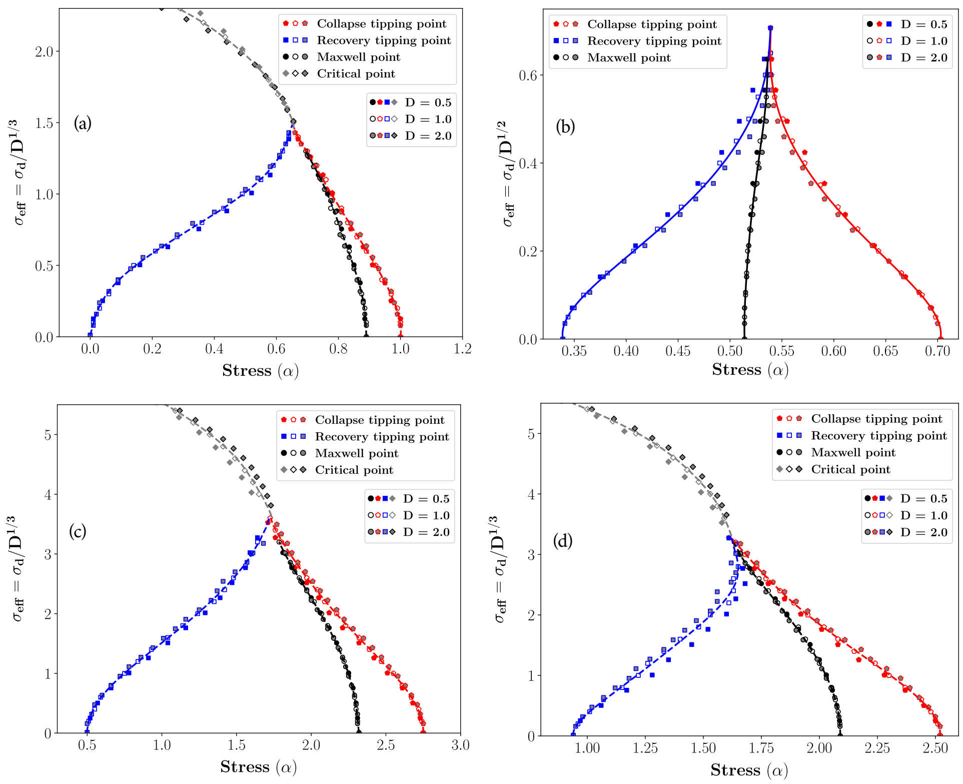


FIG. S43. **Phase diagrams in the $(\alpha, \sigma_{d,\text{eff}})$ plane for (a) desertification, (b) lake pollution, (c) grazing, and (d) insect outbreak models**. For each model we consider diffusion $D = 1/2, 1, 2$ and the curves within each plot are obtained by averaging the data collapsed across these $D$ values. We define $\sigma_{\text{eff}} = \sigma_d/D^{1/3}$ for (a), (c), and (d), and $\sigma_{d,\text{eff}} = \sigma_d/D^{1/2}$ for (b). All simulations use a system size $L \times L = 256^2$; model parameters appear in Table T1.

By and large, stochasticity drives the system towards its low-abundance sector. The overall strength of stochasticity is proportional to the size of local population, and thus it weakens in the low population sector. Therefore, the dynamics becomes slower and the system tends to get stuck in its low-abundance state [22, 23]. Consequently, in FIG.S43(a, c) the transition lines generally shift leftward: bare soil is increasingly favored as stochasticity grows. The notable exception is the recovery tipping-point line, which shifts in the opposite direction. This happens because, in the regime where vegetation invades desert, small patches still shrink while only sufficiently large patches spread [8]. As stochasticity increases, spatial fluctuations in cover raise the probability of nucleating large patches, thereby facilitating recovery.

### S5: Role of Environmental stochasticity

In this section, we investigate in detail the role of environmental stochasticity in shaping alternative stable states and the corresponding collapse and recovery tipping points. To isolate this effect, we consider dynamics that include the environmental noise and spatial dispersal in addition to the deterministic contributions, i.e.,

$$\frac{dB(\mathbf{x},t)}{dt} = \text{Deterministic terms} + \underbrace{\sigma_e B(\mathbf{x},t)\ \eta_e(\mathbf{x},t)}_{\text{Environmental stochasticity}} + \underbrace{D\ \nabla^2 B(\mathbf{x},t)}_{\text{Spatial dispersal}}. \tag{S51}$$

In parallel with Section S4, we first demonstrate how environmental stochasticity, characterized by its strength $\sigma_e$, deforms the hysteresis loops obtained via the parameter-sweep approach (Subsection J). Next, we construct phase diagrams in the $(\alpha, \sigma_e)$ parameter space using the marginal dynamics identification framework (Subsection K), and show how increasing environmental noise shifts and reshapes the various transition points. Within the same subsection, we provide a qualitative comparison of the resulting phase diagrams across different models, emphasizing both common features and model-specific differences. We also point out the agreement between the outcomes of the parameter-sweep and marginal dynamics identification methods. Finally, in Subsection L, we analyze how diffusion influences environmental stochasticity and derive an effective noise strength $\sigma_{e,\text{eff}}$ that incorporates the combined effects of $\sigma_e$ and the diffusion coefficient $D$.

##### J. Hysteresis Diagrams: Parameter sweep method

In this subsection, we present hysteresis diagrams for the four models at progressively higher levels of environmental stochasticity ($\sigma_e$), demonstrating that increasing $\sigma_e$ gradually transforms a sharp, effectively irreversible transition into a smooth and reversible one (see Fig. S51).

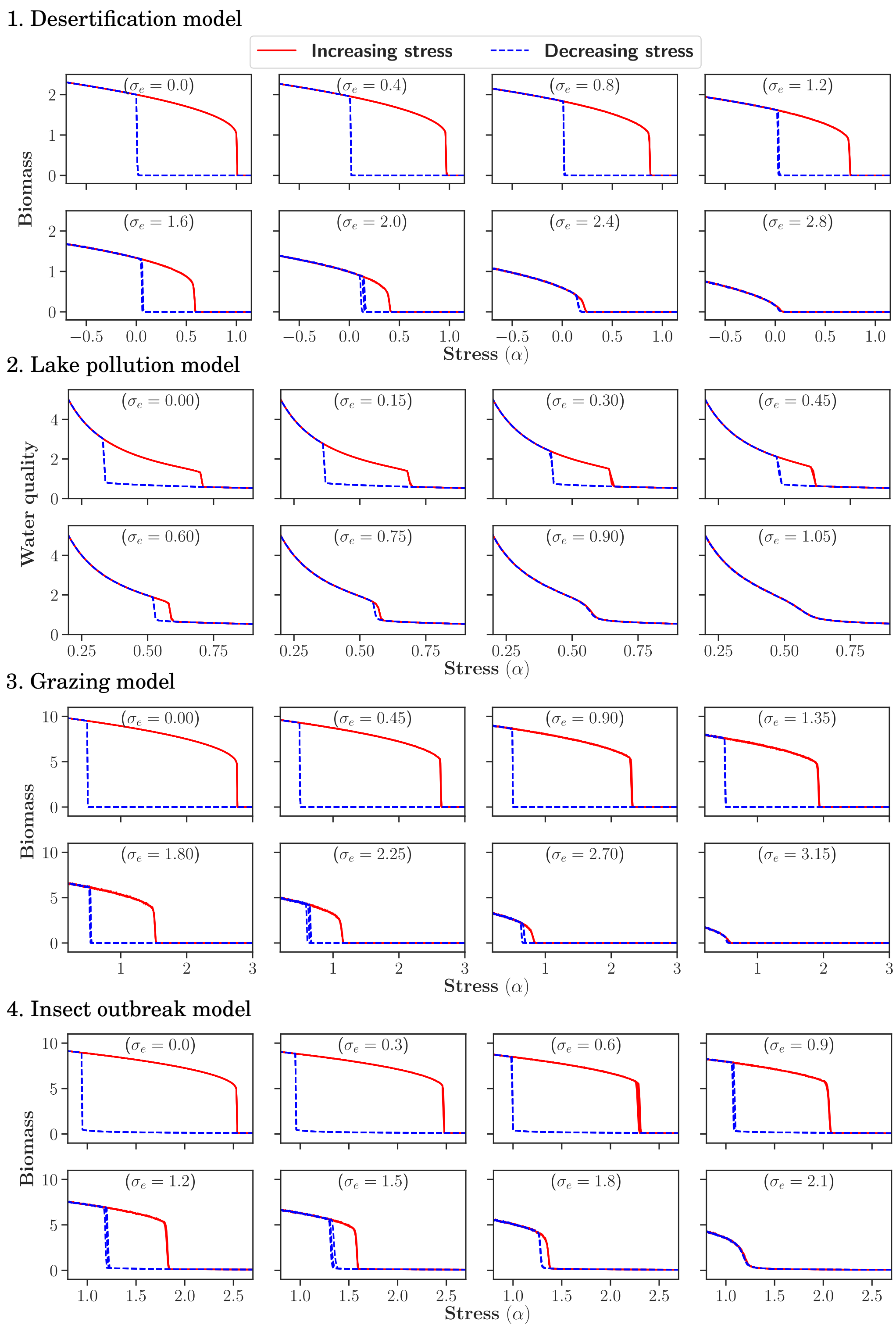


FIG. S51. Hysteresis plots for the 1. desertification, 2. lake pollution, 3. grazing, and 4. insect outbreak models. For each model, we show eight diagrams corresponding to increasing levels of environmental stochasticity, with four realizations for each diagram. Model parameters are listed in Table T1; additionally, we use dispersal rate $D = 1$, sweep rate $\epsilon = 10^{-5}$, and system size $L \times L = 256^2$.

##### K. Phase Diagrams: Marginal dynamics identification

Here we present complete phase diagrams in $(\alpha - \sigma_e)$ plane, obtained through marginal dynamics identification for all four models (see FIG. S52). For the desertification and grazing models (FIG. S52(a) and (c)), we identify the Maxwell point, the collapse and recovery tipping points, as well as an additional critical point.

Technically speaking, the dynamic of Eq.( S51) does not allow for an absorbing state. The $\sigma_e$ term makes the linear growth rate random, so the system may recover from very low densities through many good years, i.e., through an improbable sequence of high growth rates. Still (see explanation below), the strong-noise transition for desertification and grazing models admits a critical behavior with the exponents of the directed percolation transition. In contrast, the lake pollution and insect outbreak models (FIG. S52(b) and (d)) do not exhibit an absorbing state, as both stationary state solutions are non-zero. Consequently, no critical point appears in the strong-stochasticity sector of the phase diagrams for these models. Notably, unlike demographic stochasticity in the insect outbreak model (FIG. S42(c)), environmental stochasticity cannot transform the lowest non-zero stationary state into an absorbing state.

All phase diagrams additionally display tipping points obtained from the parameter-sweep approach, indicated by dotted and dashed curves in each panel. For every model, these tipping points are shown for two distinct sweep rates, $\epsilon = 10^{-5}$ and $10^{-4}$, illustrating the complementary nature of the two methodologies. As the sweep rate $\epsilon$ is reduced, the tipping points identified via parameter sweeping progressively approach those determined using marginal dynamics identification.

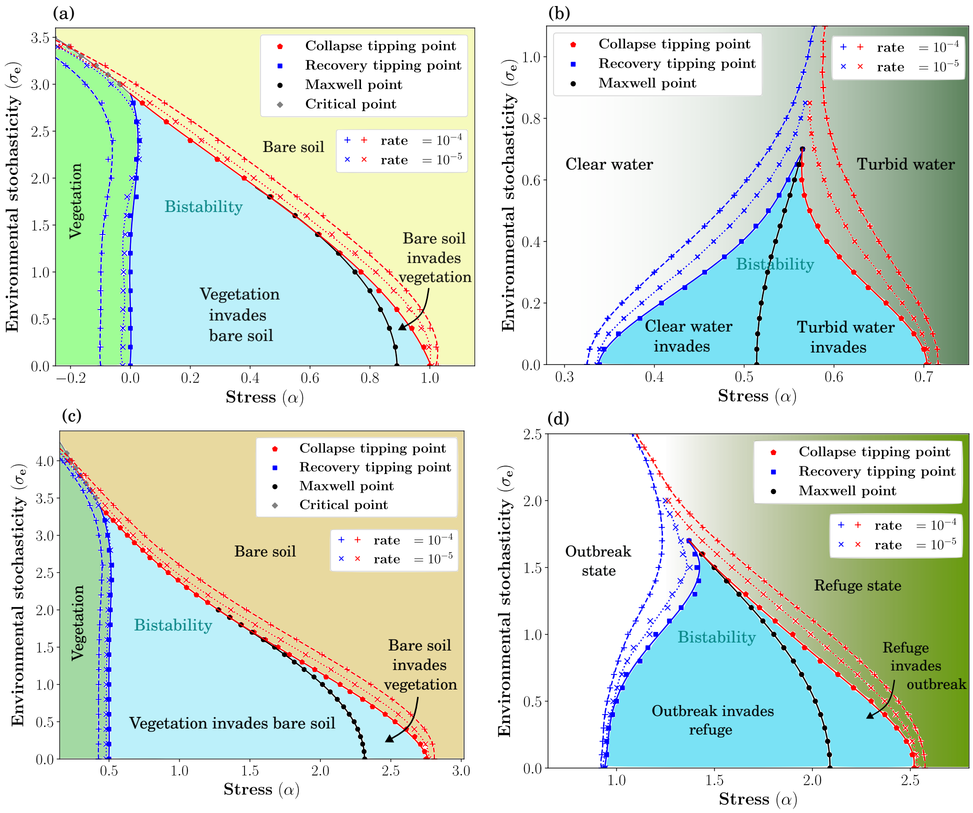


FIG. S52. **Phase diagrams in the $(\alpha - \sigma_e)$ plane for (a) desertification, (b) lake pollution, (c) grazing, and (d) insect outbreak models.** Solid lines denote phase boundaries obtained using the marginal dynamics identification method. Regions shaded in light cyan correspond to bistable domains where one of the stationary states is absorbing, whereas dark cyan regions indicate bistability between two non-zero stationary states. Dotted ($\times$) and dashed ($+$) lines show tipping-point trajectories obtained from the parameter-sweep method for rates $\epsilon = 10^{-5}$ and $\epsilon = 10^{-4}$, respectively. Model parameters are listed in Table T1 with additional dispersal rate $D = 1$ and system size $L \times L = 256^2$.

#### L. Effect of diffusion on environmental stochasticity

Following a similar argument used for $\sigma_d$ and $D$ (see Supplementary I), we define an effective stochasticity $\sigma_{e,\text{eff}}$ that combines $\sigma_e$ and $D$. From numerical simulations, we find $\sigma_{e,\text{eff}} = \sigma_e/D^{\zeta}$. For models with two nonzero stationary solutions at the deterministic limit, such as the lake pollution and insect outbreak models, setting $\zeta = 1/2$ produces a clear data collapse within the bistable regime (see FIG.S53(b),(d)). For models with a zero stationary solution at the deterministic limit, such as the desertification and grazing models, $\zeta = 1/3$ yields a consistent collapse (see FIG.S53(a),(c)).

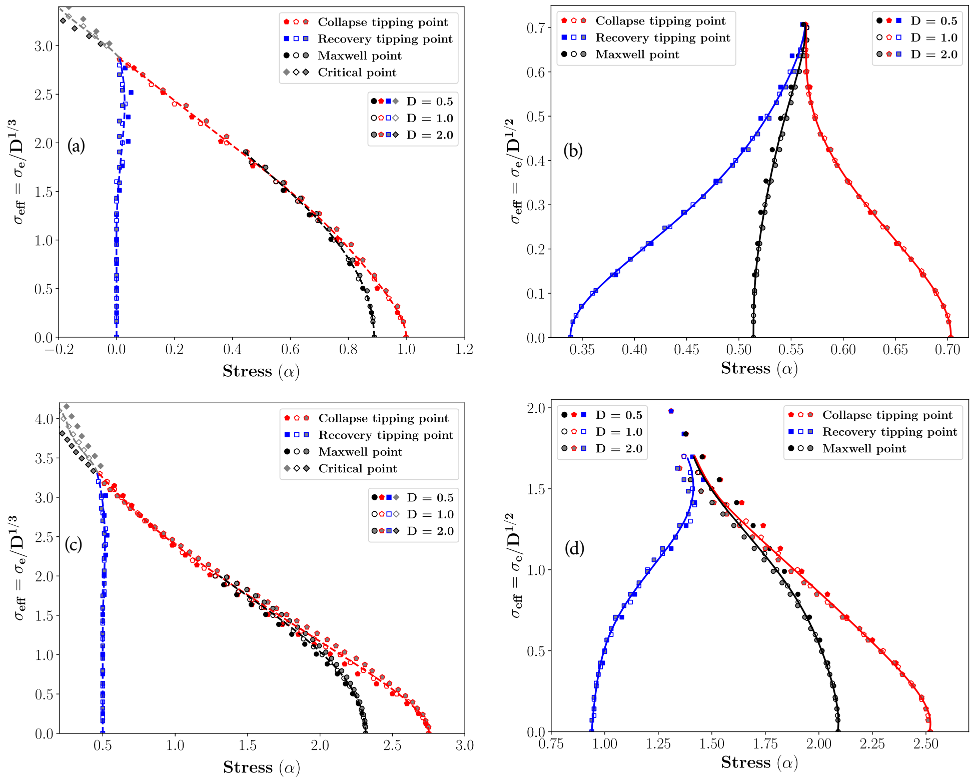


FIG. S53. **Phase diagrams in the** $(\alpha, \sigma_{e,\text{eff}})$ **plane for (a) desertification, (b) lake pollution, (c) grazing, and (d) insect outbreak models**. For each model we consider diffusion $D = 1/2, 1, 2$ and the curves within each plot are obtained by averaging the data collapsed across these $D$ values. We define $\sigma_{e,\text{eff}} = \sigma_e/D^{1/3}$ for (a) and (c), and $\sigma_{e,\text{eff}} = \sigma_e/D^{1/2}$ for (b) and (d). All simulations use a system size $L \times L = 256^2$; model parameters appear in Table T1.

#### M. Inclination of the recovery tipping point

From FIG. S42(a,c) and FIG. S52(a,c), it is evident that the recovery tipping point shows (almost) no inclination under environmental stochasticity, while under demographic stochasticity it is inclined to the right. In other words, strong demographic stochasticity generates strong abundance fluctuations, leading to immediate nucleation and expansion of the healthy state, whereas environmental stochasticity fails to generate such strong fluctuations.

To understand better this behavior let us consider a simple, single-patch model for dilute vegetation, so only the linear terms are relevant. The corresponding Fokker-Planck equation for demographic stochasticity in the desertification model is,

$$\frac{\partial P(B,t)}{\partial t} = \alpha \frac{\partial}{\partial B}[BP(B,t)] + \frac{\sigma_d^2}{2}\frac{\partial^2}{\partial B^2}[BP(B,t)] - \mu \frac{\partial}{\partial B} P(B,t). \tag{S52}$$

where $\mu$ is the migration rate. The steady state solution of this equation is,

$$P_{ss}(B) = A_d \frac{e^{-2\alpha B/\sigma_d^2}}{B^{1-2\mu/\sigma_d^2}}. \tag{S53}$$

where $A_d$ is a normalization factor.

The corresponding Fokker-Planck equation for environmental stochasticity is,

$$\frac{\partial P(B,t)}{\partial t} = \alpha \frac{\partial}{\partial B}[BP(B,t)] + \frac{\sigma_e^2}{2}\frac{\partial^2}{\partial B^2}[B^2 P(B,t)] - \mu \frac{\partial}{\partial B} P(B,t). \tag{S54}$$

Now the steady state, zero flux solution of this equation is,

$$P_{\mathrm{ss}}(B) = A_e B^{-\left(2+\frac{2\alpha}{\sigma_e^2}\right)} \exp\left(-\frac{2\mu}{\sigma_e^2 B}\right) \tag{S55}$$

where $A_e$ is a normalization factor.

Comparing the distribution (S53) with (S55) for small values of $\mu$ (we introduced $\mu$ only to avoid absorption) and $\alpha$, one finds that under demographic stochasticity the large-$B$ behavior decreases asymptotically as $1/B$, whereas under environmental stochasticity it decreases like $1/B^2$. Therefore, large fluctuations are much rarer under environmental stochasticity. This reflects the fact that demographic stochasticity is proportional to $\sqrt{B}$, which is much larger than $B$ for dilute populations. In other words, once we put aside the extinction state at $B = 0$ itself and consider only the dilute state $B \to 0$, the stickiness (*sensu* [22, 23]) under demographic stochasticity is weaker than the stickiness under environmental stochasticity. A similar line of argument holds for the grazing model.

#### S6: Role of Spatial heterogeneity

Below we examine in detail how spatial heterogeneity shapes alternative stable states and the associated collapse and recovery tipping points. To focus exclusively on this effect, we consider dynamics that include spatial heterogeneity and dispersal in addition to the deterministic contributions, i.e.,

$$\frac{dB(\mathbf{x},t)}{dt} = \text{Deterministic components with } \alpha \text{ replaced by } \underbrace{\alpha(\mathbf{x}) \to \alpha + \Delta\xi(\mathbf{x})}_{\text{Spatial heterogeneity}} + \underbrace{D\,\nabla^2 B(\mathbf{x},t)}_{\text{Spatial dispersal}}.$$

Following the presentation in the last sections, we first show how spatial heterogeneity, controlled by $\Delta$, modifies the hysteresis loops obtained using the parameter-sweep approach (Subsection N).

With spatial heterogeneity, the transition may become continuous but irreversible, therefore we were looking for a parameter that will characterize a transition between abrupt-and-irreversible (catastrophic) shift to continuous-and-irreversible shift. In Subsection O, we define such an abruptness parameter for spatially heterogeneous systems with dispersal. Armed with this abruptness parameter we construct, in Subsection P, abruptness heatmaps in the $(D, \Delta)$ plane. Finally, in Subsection Q we discuss how diffusion modulates the influence of heterogeneity.

##### N. Hysteresis Diagrams: Parameter sweep method

In this subsection, we present hysteresis diagrams for the four models at progressively higher levels of spatial heterogeneity ($\Delta$). As with environmental and demographic stochasticity, spatial heterogeneity tends to reduce or eliminate hysteresis (see Fig. S61). However, whereas stochasticity-induced hysteresis typically involves an abrupt jump in the system's state, spatial heterogeneity can produce hysteretic (irreversible) behavior even when the system-wide transition is continuous. The results for the insect outbreak model with $\Delta = 8.0$ (Fig. S61) provide a pronounced example.

More generally, even in other cases, increasing the resolution in $\alpha$ can reveal that what initially appears to be a sharp jump is in fact a continuous transition. Therefore, we require a more quantitative measure of the transition's abruptness.

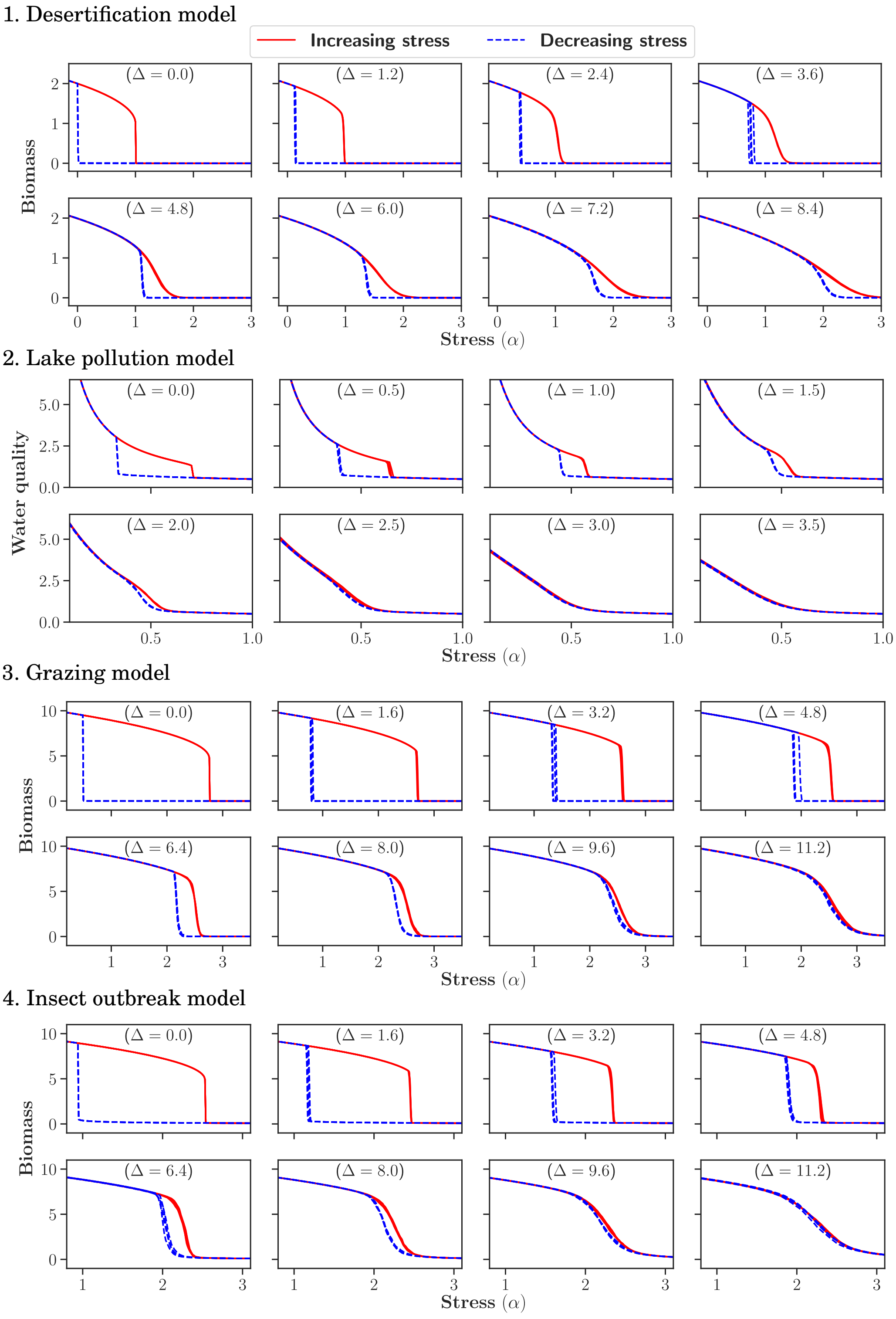


FIG. S61. Hysteresis plots for the 1. desertification, 2. lake pollution, 3. grazing, and 4. insect outbreak models. For each model, we show eight diagrams corresponding to increasing levels of spatial heterogeneity, with four realizations for each diagram. Model parameters are listed in Table T1; additionally, we use dispersal rate $D = 1$, sweep rate $\epsilon = 10^{-5}$, and system size $L \times L = 256^2$.

##### O. The abruptness parameter: characterizing the sharpness of irreversible transitions in spatially heterogeneous systems with dispersal

It is known that when spatial heterogeneity is low, ecosystems respond coherently to external stress, making them prone to abrupt and catastrophic transitions. Under strong spatial heterogeneity, spatial dispersal plays an important role by mediating the impact of heterogeneity on ecosystem dynamics [24]. Low dispersal allows local patches to respond independently, preserving spatial heterogeneity and reducing the likelihood of synchronized collapse (FIG. S62b). In contrast, high dispersal strongly couples patches and can synchronize their dynamics, allowing regime shifts to propagate across the system and resulting in abrupt, system-wide transitions even under strong, randomly distributed heterogeneity (FIG. S62c). Thus, dispersal alters the abruptness of a transition depending on its strength relative to spatial heterogeneity.

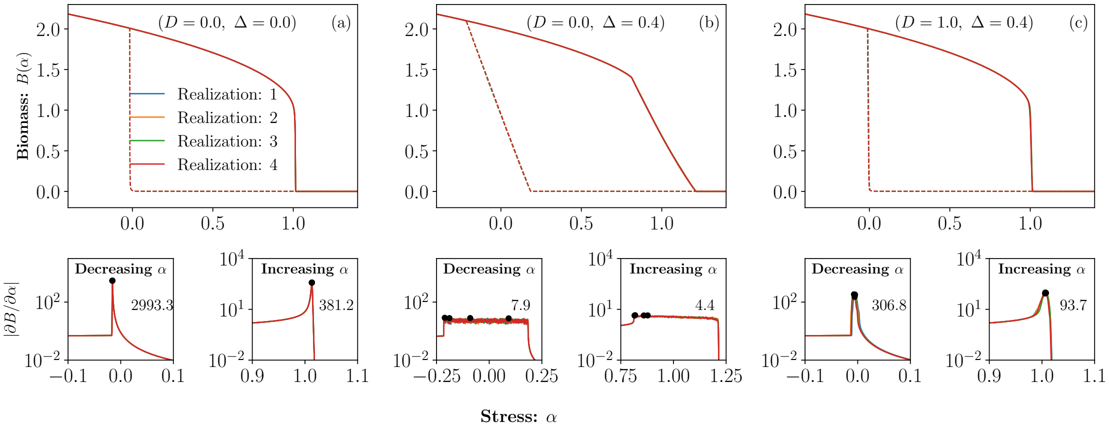


FIG. S62. Hysteresis plots for the desertification model ($\beta = 2, \gamma = 1$, with sweep rate $\epsilon = 4 \times 10^{-5}$ and system size $L \times L = 128^2$) are shown for three different sets of $[D, \Delta]$: [0, 0], [0, 0.4], and [1.0, 0.4]. In the first row we show the results of four realizations in each diagram. These four hysteresis loops are indistinguishable, indicating that (at least for these values of $\Delta$ and system sizes) the differences between different realizations are tiny. In (a) there is no heterogeneity, thus all patches collapse and recover at the same value of stress parameter. The effect of spatial heterogeneity in making the transition gradual is shown in (b); there is no dispersal, and every patch switches to the alternative state at a different value of $\alpha$. The interplay between dispersal and heterogeneity is demonstrated in panel (c): an increase in dispersal makes the transition more abrupt. For each hysteresis diagram in the first row, $|\partial B/\partial \alpha|$ is plotted directly below in the second row. The abruptness parameter, i.e., the maximum of $|\partial B/\partial \alpha|$, is indicated by filled black circles, with the corresponding average value reported within each panel.

To quantify the abruptness of a transition, that is, how sharply the biomass changes as the stress parameter $\alpha$ is varied in the presence of spatial heterogeneity ($\Delta$) and dispersal ($D$), we introduce our **abruptness parameter**. We define this quantity as the maximum magnitude of the change of the mean system's biomass with respect to the stress parameter $\alpha$, i.e., $\max |\partial B/\partial \alpha|$, evaluated along increasing or decreasing sweeps of $\alpha$ at a constant pace (see below for further discussion of the abruptness parameter).

The rationale goes as follows. In homogeneous systems, ($\Delta = 0$) the abrupt jump is global, i.e., it involves all the patches, so the discontinuous jump in biomass density, as in FIG. S62(a), is large. Correspondingly, our abruptness parameter takes its maximal value. As $\Delta$ increase this discontinuous jump becomes smaller in magnitude as in FIG. S62(c), and finally it disappears completely. In what follows, and in the main text, we normalize the abruptness parameter by its $\Delta = 0$ value, so it indicates how severe is the catastrophic shift in units of its maximal effect.

#### P. Methodology for constructing heatmaps of abruptness parameter in $(D, \Delta)$ plane

To construct a heatmap of abruptness parameter in the $(D, \Delta)$ plane, we consider a dense grid over a suitable range of $D$ and $\Delta$ (see FIG. S63a). To improve accuracy, we use the average of 16 independent realizations, followed by Gaussian smoothing along both axes (FIG. S63b). As explained, we normalize the abruptness parameter by its maximum value at $\Delta = 0$, so all the results are in the interval $[0, 1]$. For clarity, we overlay contours (isoclines) at fixed levels 0.1–0.9, as shown in FIG. S63c.

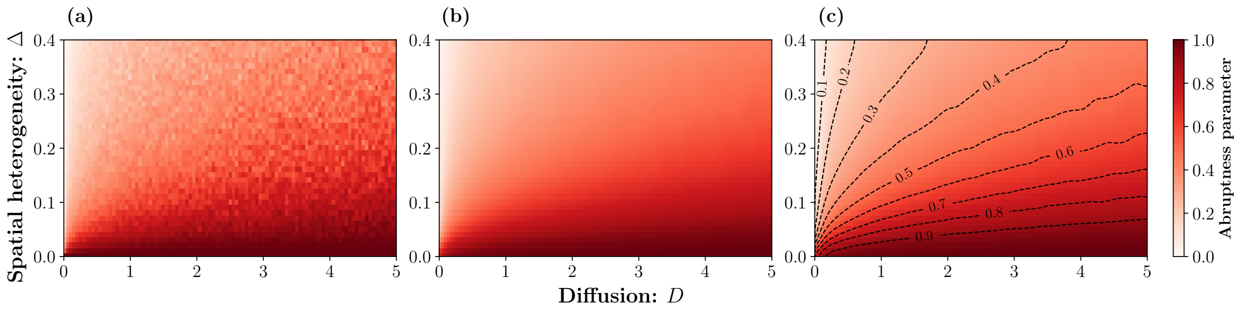


FIG. S63. Heatmap of abruptness parameter in the $(D, \Delta)$ plane for the collapse tipping point of the desertification model ($\beta = 2, \gamma = 1$, $L = 128$ with sweep rate $\epsilon = 4 \times 10^{-5}$). (a) A dense $100 \times 40$ grid over $D \in [0, 5]$ and $\Delta \in [0, 0.4]$ is shown for one realization. (b) Averaging over 16 independent realizations followed by Gaussian smoothing yields a smoother surface. (c) Isoclines at fixed levels 0.1–0.9 are included alongside the heatmap after normalizing the parameter to the interval $[0, 1]$.

#### Q. Effect of diffusion on spatial heterogeneity

As manifested in FIGs S64, and as expected, the severity of the catastrophic shift, i.e., the abruptness parameter, decreases monotonically as $\Delta$ increases (due to fragmentation of the system, where each fragment switches to the alternative state independently) and decreases as $D$ increases, since diffusion wipes out the effect of spatial heterogeneity. Overall, the effect is governed by the combined parameter $\Delta^2/D$, as discussed in the main text.

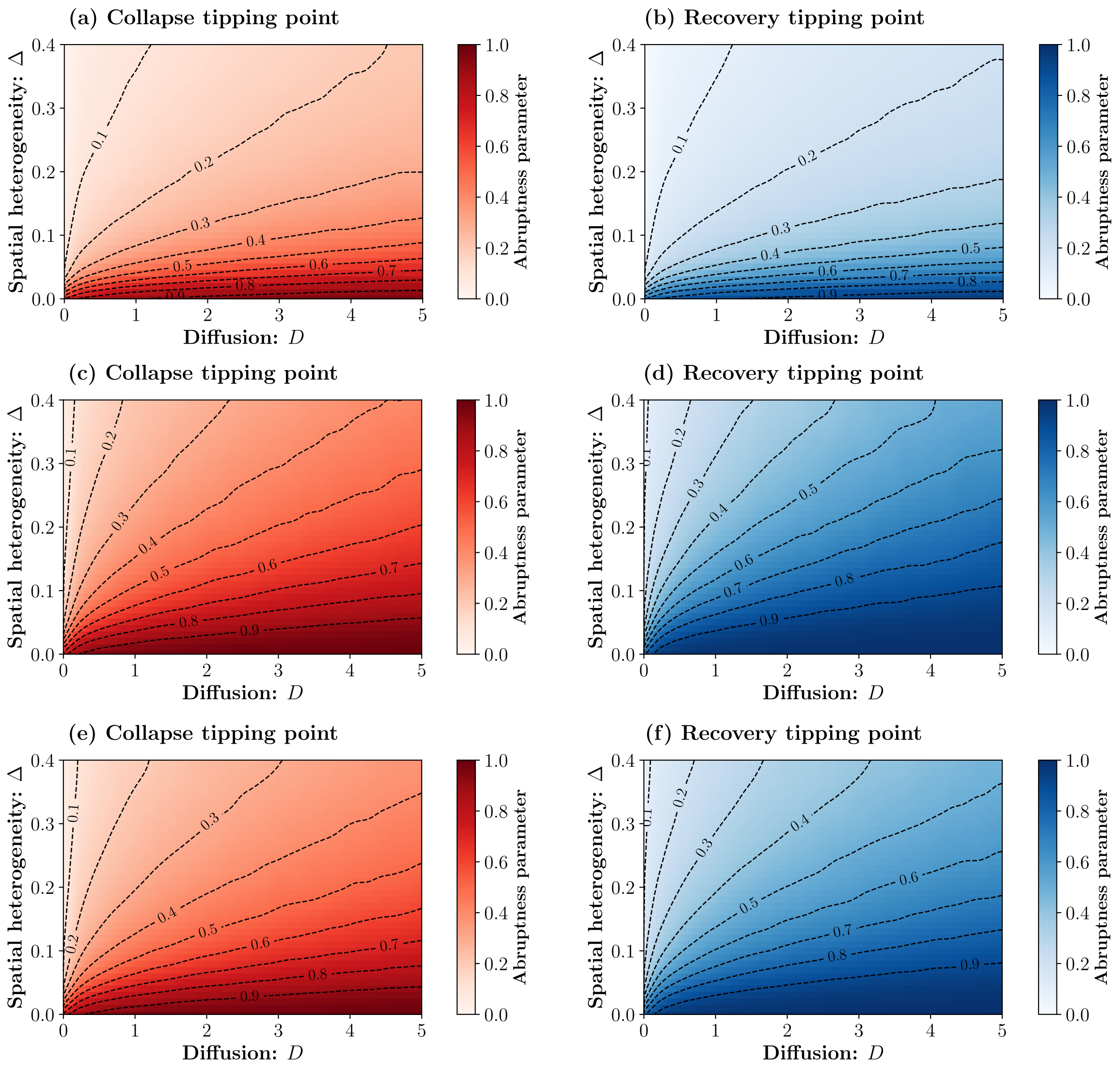


FIG. S64. **Heatmap of abruptness parameter in the $(D, \Delta)$ plane:** Normalized abruptness parameter for the collapse and recovery tipping points are shown along the first and second columns, respectively, for the (a, b) lake pollution, (c, d) grazing, and (e, f) insect outbreak models. In all cases, the sweep rate $\epsilon = 4 \times 10^{-5}$ and system size $L \times L = 128^2$, with the remaining model parameters listed in Table T1.

##### R. Heterogeneous systems and the abruptness parameter.

Throughout this section, we consider a heterogeneous system composed of many components (patches). Each component may undergo a discontinuous and irreversible transition at a distinct value of the external parameter (stress), while a concurrent "smoothing" process (migration) acts to reduce differences among their states. The interplay between heterogeneity and this smoothing process governs the dynamics of the shift.

The same problem arises in other fields, in particular in the zero-temperature dynamics of heterogeneous systems that exhibit first-order transitions, such as the random-field Ising model [25]. Here we adopted a more "ecological" approach, so our abruptness parameter is defined in terms of quantities (such as total biomass) that are both measurable and of direct interest, under reasonable conditions (e.g., system size, sweep rate, etc.). We conclude this section by commenting on several issues related to our approach.

- **System's size and rare events:** In heterogeneous systems rare events may dominate the dynamics [26]. These rare events have to do with exponentially rare spatial configurations of $\alpha(\mathbf{x})$, say, that has a large effect on the outcomes. When these effects are important, the sample-to-sample fluctuations are huge.

  We have chosen values of $L$ and $\Delta$ that are, on the one hand, comparable with what one expects to find in the field and, on the other hand, small enough so rare spatial configurations do not appear. Therefore, different realization yield very similar results. Moreover, we checked that our abruptness parameter converges to a given value as system size increases in the range considered here (up to $L = 160$), and is, more or less, independent of the sweep rate.

- **The transition:** The physics literature (e.g., [25]) typically focuses on whether a jump is macroscopic or microscopic. For example, in the random-field Ising model, weak heterogeneity yields a single jump in which the number of spins that flip constitutes a finite fraction of the total number of spins in the system, whereas under strong heterogeneity one observes only microscopic jumps.

  While this approach is mathematically rigorous, its implementation requires a sweep rate that is much slower than the timescale associated with the magnetic dynamics. This requirement is reasonable in magnetic systems, but it is unrealistic in ecological settings, where the system's response is comparatively slow. Accordingly, we introduced our abruptness parameter. As implemented here (with a finite sweep rate), the goal of this parameter is not to distinguish transition types in a rigorous mathematical sense, but rather to quantify how large a jump one should expect under different conditions.

### S7: Additivity: the combined effects of stochasticity and spatial heterogeneity

Let us consider, now, how different complexities combine and influence the phase diagram. As stated in the main text, our main conclusion is that, at least for in a substantial range of the parameter space, the effect is additive, i.e., one may consider each of these factors separately and add the results.

In what follows we provide a detailed analysis of the combined effects. First, in Subsection S, we consider the three pairwise combinations, $(\sigma_d, \sigma_e)$, $(\sigma_d, \Delta)$, and $(\sigma_e, \Delta)$, and systematically analyze the hysteresis diagrams obtained using the parameter-sweep method. Subsection T then outlines the methodology for constructing heatmaps of these indices in the corresponding parameter planes. Finally, in Subsection U, we present these heatmaps for each of the three parameter combinations across the different models.

#### S. Hysteresis diagrams by the parameter-sweep method

In this subsection, we systematically analyze hysteresis diagrams obtained using the parameter-sweep method across all four models, considering three pairwise combinations of complexities. For each model, we plot hysteresis diagrams for the three parameter pairings $(\sigma_d, \sigma_e)$, $(\sigma_d, \Delta)$, and $(\sigma_e, \Delta)$, yielding twelve cases in total.

In each case, we present 25 hysteresis diagrams arranged in a $5 \times 5$ grid, corresponding to different parameter combinations. Each diagram includes 4 realizations to illustrate stochastic variability. For example, for the $(\sigma_d, \sigma_e)$ pairing in the desertification model (see Fig. S71), we plot 25 diagrams in which each row corresponds to increasing $\sigma_d$ at fixed $\sigma_e$, and each column corresponds to increasing $\sigma_e$ at fixed $\sigma_d$. The first row therefore isolates the effect of $\sigma_d$ on hysteresis, while the first column isolates the effect of $\sigma_e$. The remaining grid entries capture their combined influence. This layout makes it possible to track how increasing either or both $\sigma_d$

and $\sigma_e$ alters degradation, the restoration gap, and the hysteresis area, and how transitions shift from abrupt and effectively irreversible to smoother and more reversible behavior.

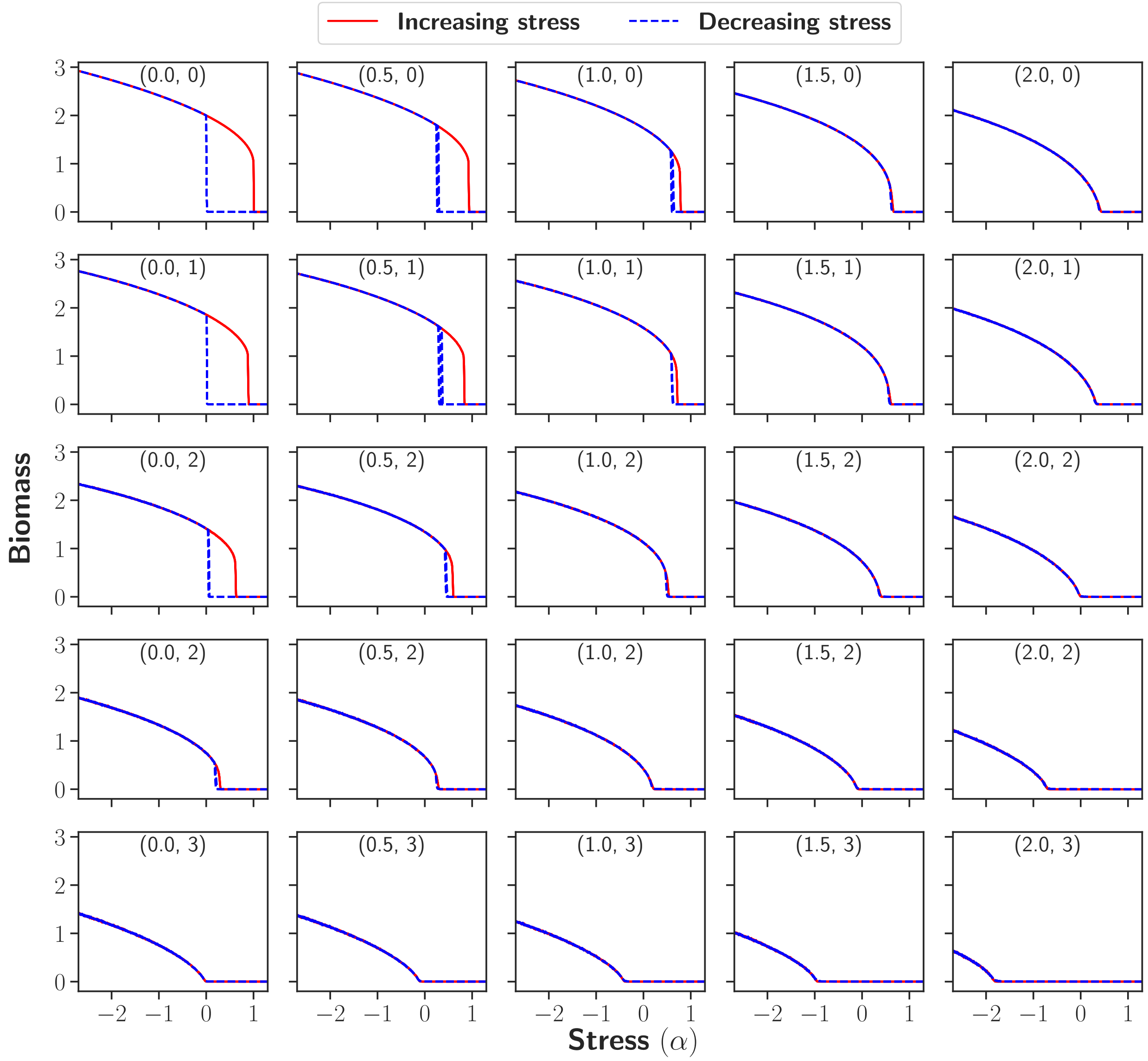


FIG. S71. Hysteresis diagrams illustrating the effects of different $(\sigma_d, \sigma_e)$ combinations in the desertification model. Model parameters: $\beta = 2$, $\gamma = 1$, dispersal rate $D = 1$, sweep rate $\epsilon = 10^{-5}$, and the system size $L \times L = 256^2$. The relevant value of $(\sigma_d, \sigma_e)$ appears at the top of each sub-panel.

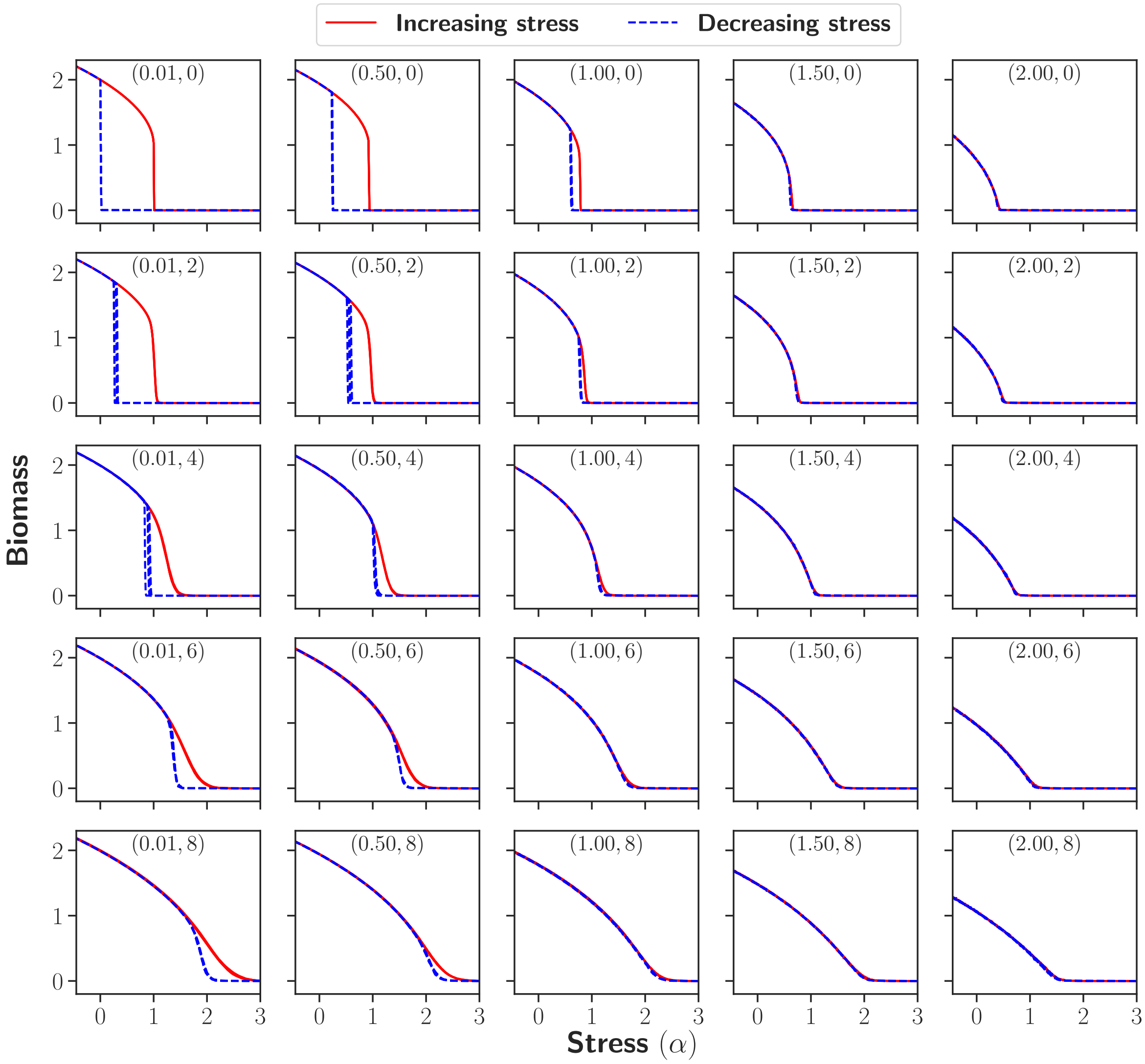


FIG. S72. Hysteresis diagrams illustrating the effects of different $(\sigma_d, \Delta)$ configurations in the desertification model. Model parameters: $\beta = 2$, $\gamma = 1$, dispersal rate $D = 1$, sweep rate $\epsilon = 10^{-5}$, and the system size $L \times L = 256^2$. The relevant value of $(\sigma_d, \Delta)$ appears at the top of each sub-panel.

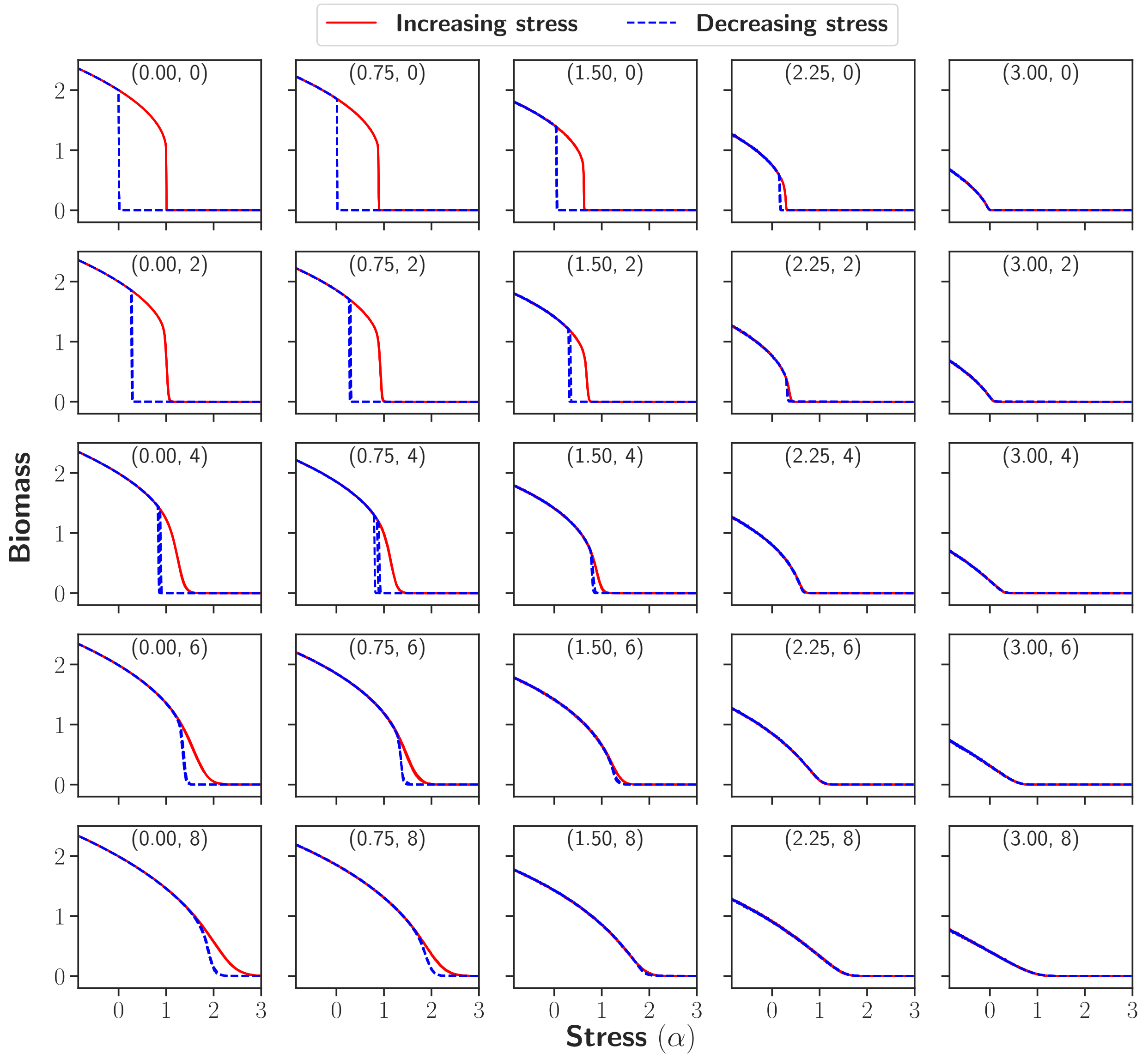


FIG. S73. Hysteresis diagrams representing the influence of changing $(\sigma_e, \Delta)$ values in the desertification model. Model parameters: $\beta = 2$, $\gamma = 1$, dispersal rate $D = 1$, sweep rate $\epsilon = 10^{-5}$, and the system size $L \times L = 256^2$. The relevant value of $(\sigma_e, \Delta)$ appears at the top of each sub-panel.

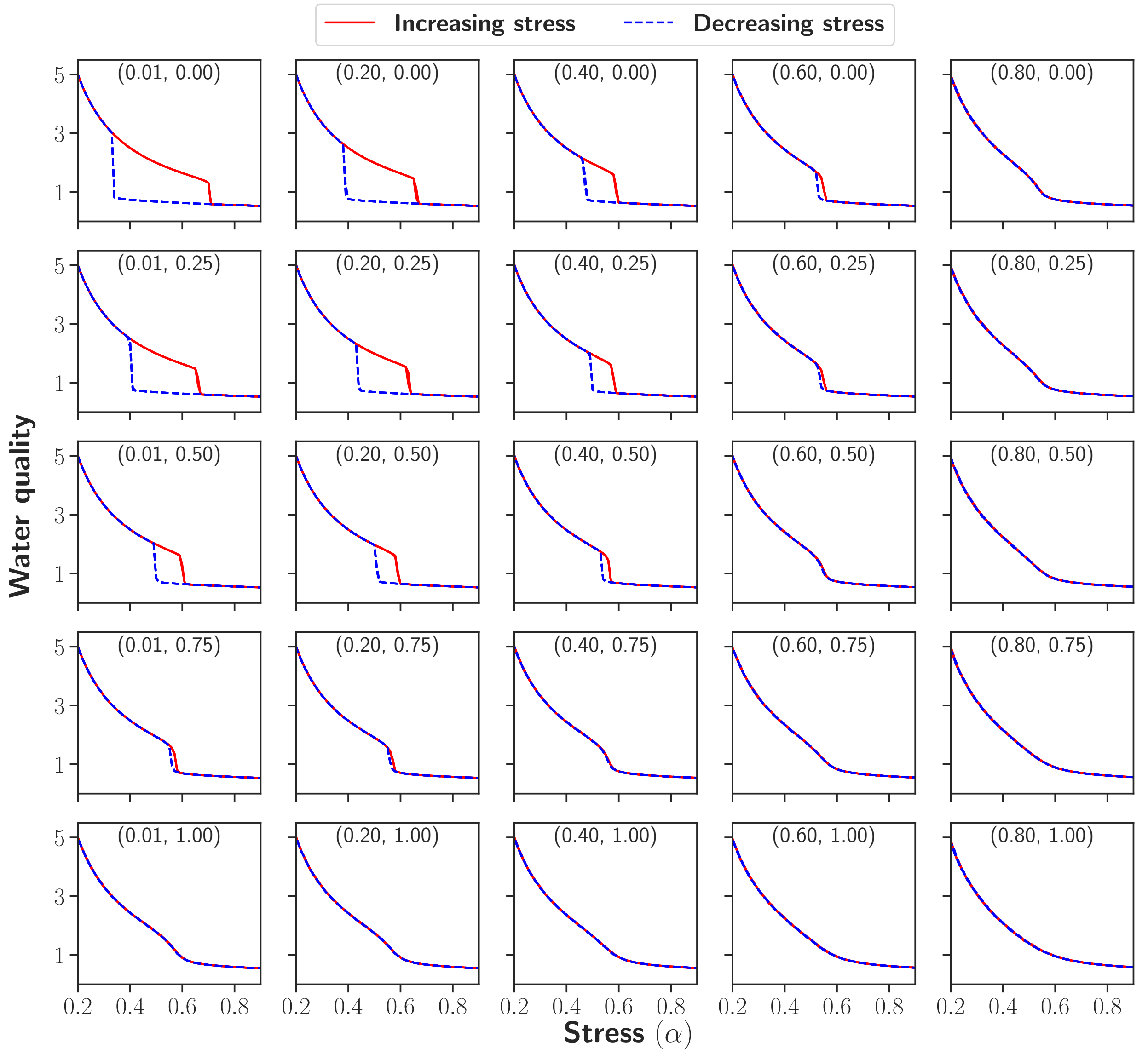


FIG. S74. Hysteresis diagrams illustrating the effects of different $(\sigma_d, \sigma_e)$ combinations in the lake pollution model. Model parameters: $\beta = r = \delta = 1$, $p = 10$, dispersal rate $D = 1$, sweep rate $\epsilon = 10^{-5}$, and the system size $L \times L = 256^2$. The relevant value of $(\sigma_d, \sigma_e)$ appears at the top of each sub-panel.

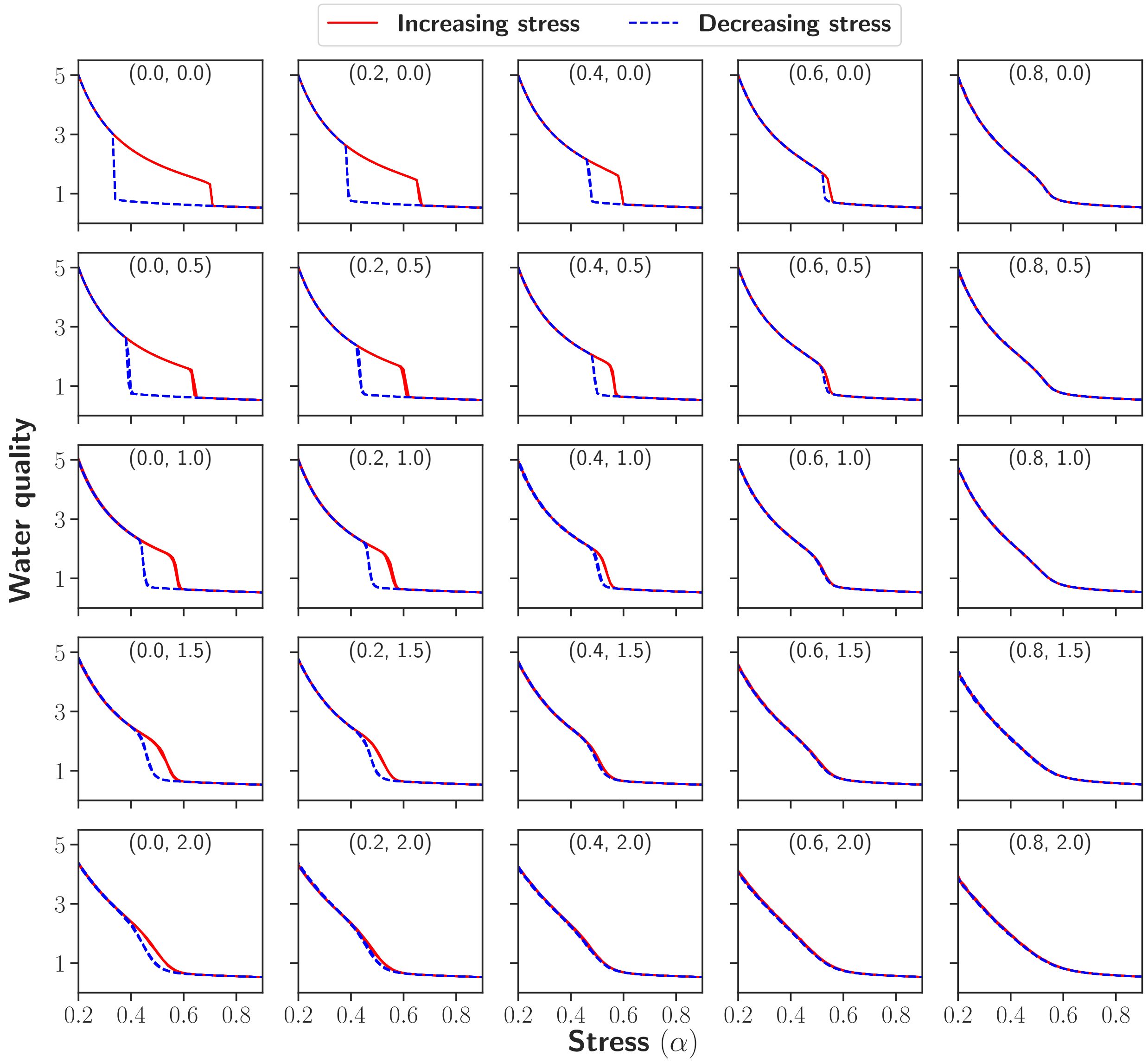


FIG. S75. Hysteresis diagrams showing the impact of various $(\sigma_d, \Delta)$ configurations in the lake pollution model. Model parameters: $\beta = r = \delta = 1$, $p = 10$, dispersal rate $D = 1$, sweep rate $\epsilon = 10^{-5}$, and the system size $L \times L = 256^2$. The relevant value of $(\sigma_d, \Delta)$ appears at the top of each sub-panel.

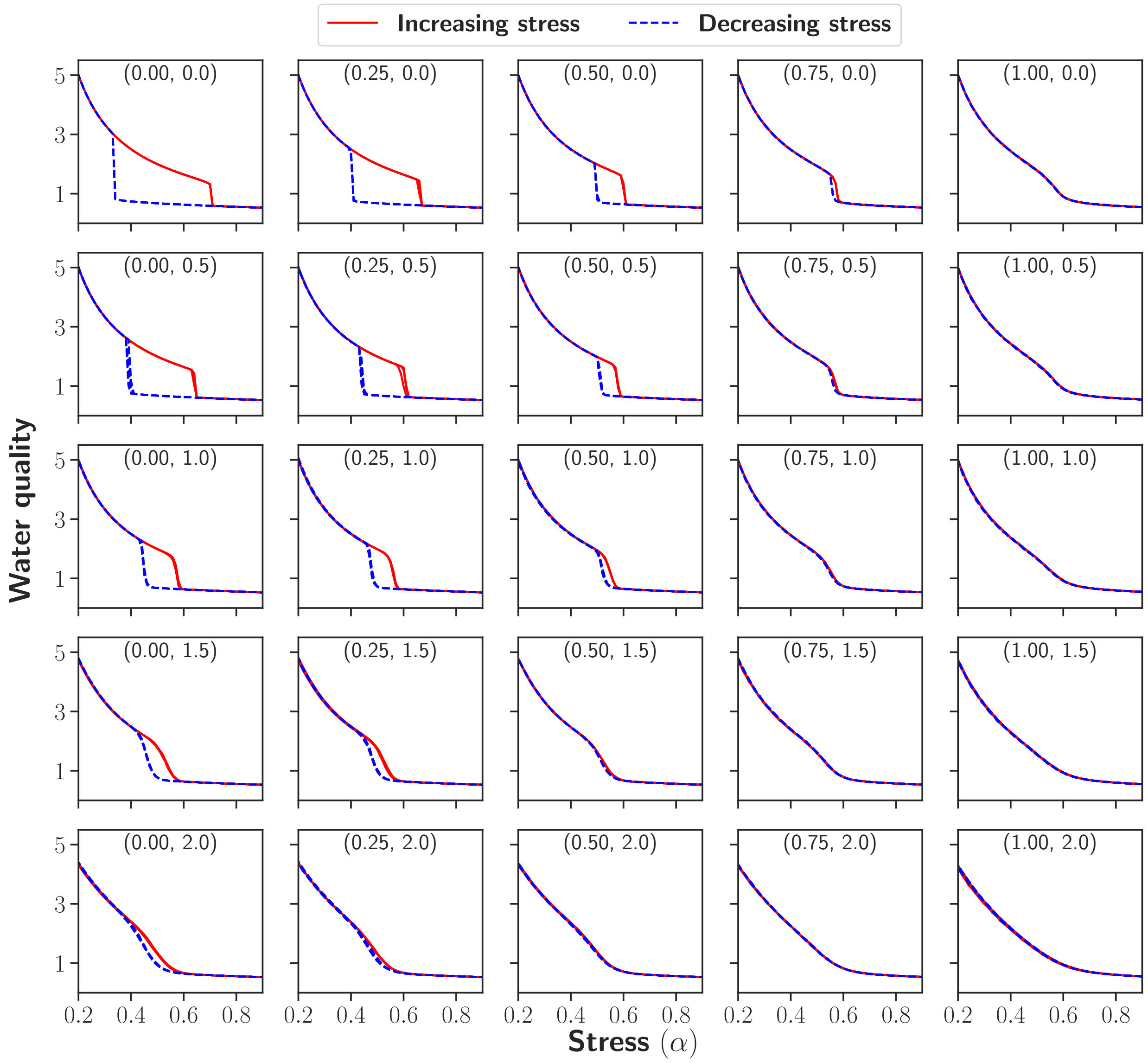


FIG. S76. Hysteresis diagrams representing the influence of changing $(\sigma_e, \Delta)$ values in the lake pollution model. Model parameters: $\beta = r = \delta = 1$, $p = 10$, dispersal rate $D = 1$, sweep rate $\epsilon = 10^{-5}$, and the system size $L \times L = 256^2$. The relevant value of $(\sigma_e, \Delta)$ appears at the top of each sub-panel.

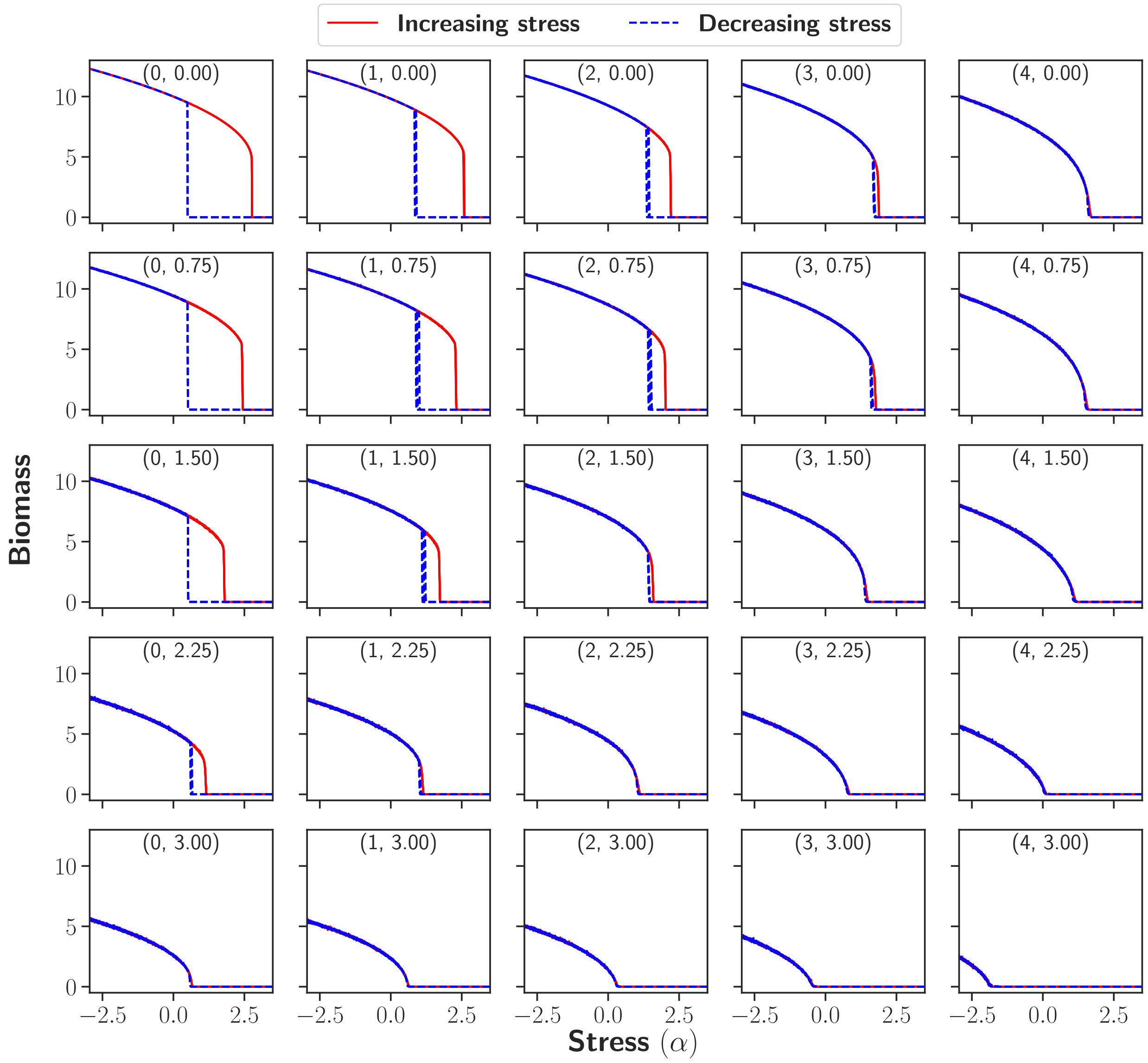


FIG. S77. Hysteresis diagrams showing the impact of various $(\sigma_d, \sigma_e)$ configurations in the grazing model. Model parameters: $r = 1$, $K = 10$, $\delta = 0.5$, dispersal rate $D = 1$, sweep rate $\epsilon = 10^{-5}$, and the system size $L \times L = 256^2$. The relevant value of $(\sigma_d, \sigma_e)$ appears at the top of each sub-panel.

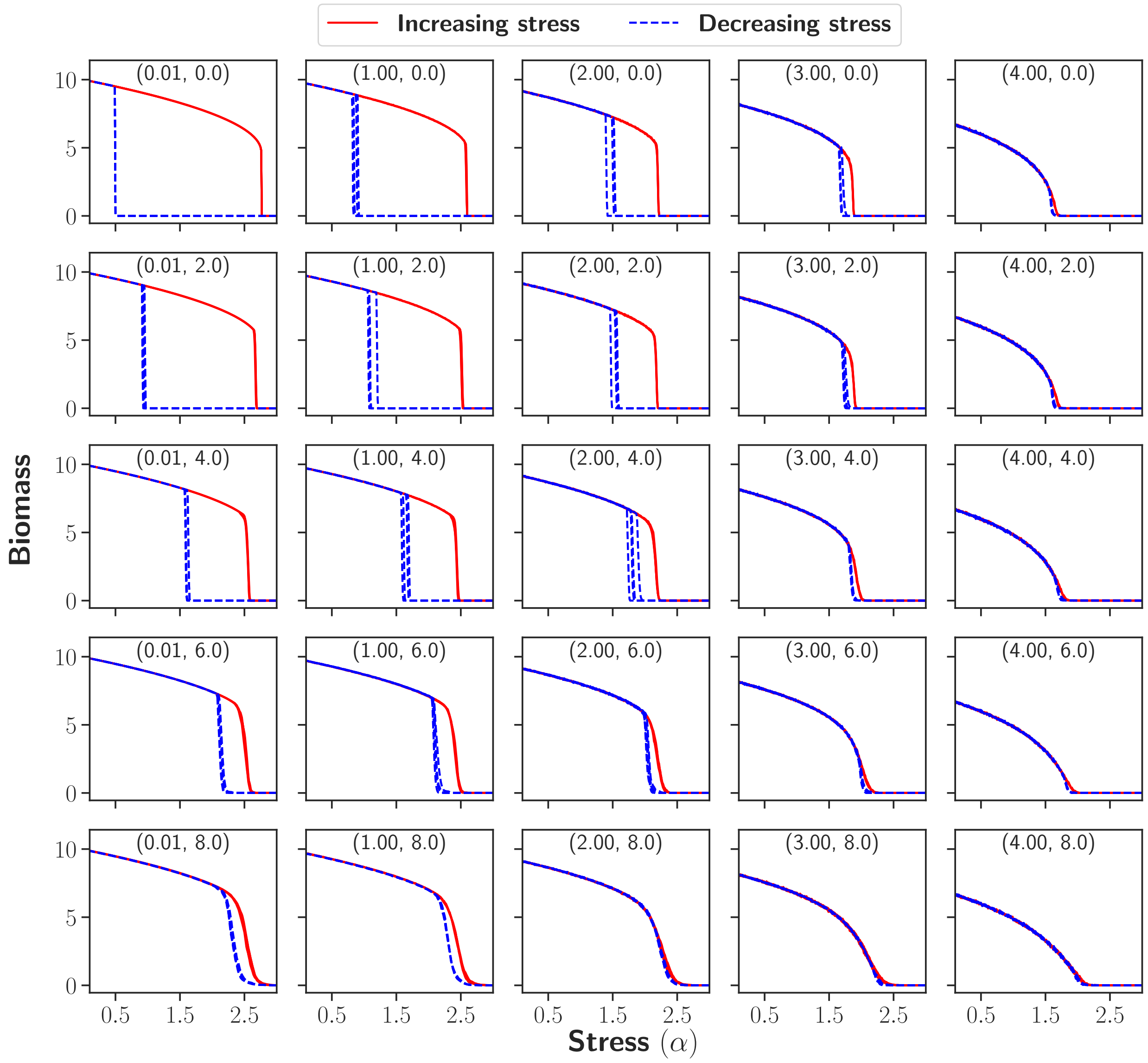


FIG. S78. Hysteresis diagrams showing the impact of various $(\sigma_d, \Delta)$ configurations in the grazing model. Model parameters: $r = 1$, $K = 10$, $\delta = 0.5$, dispersal rate $D = 1$, sweep rate $\epsilon = 10^{-5}$, and the system size $L \times L = 256^2$. The relevant value of $(\sigma_d, \Delta)$ appears at the top of each sub-panel.

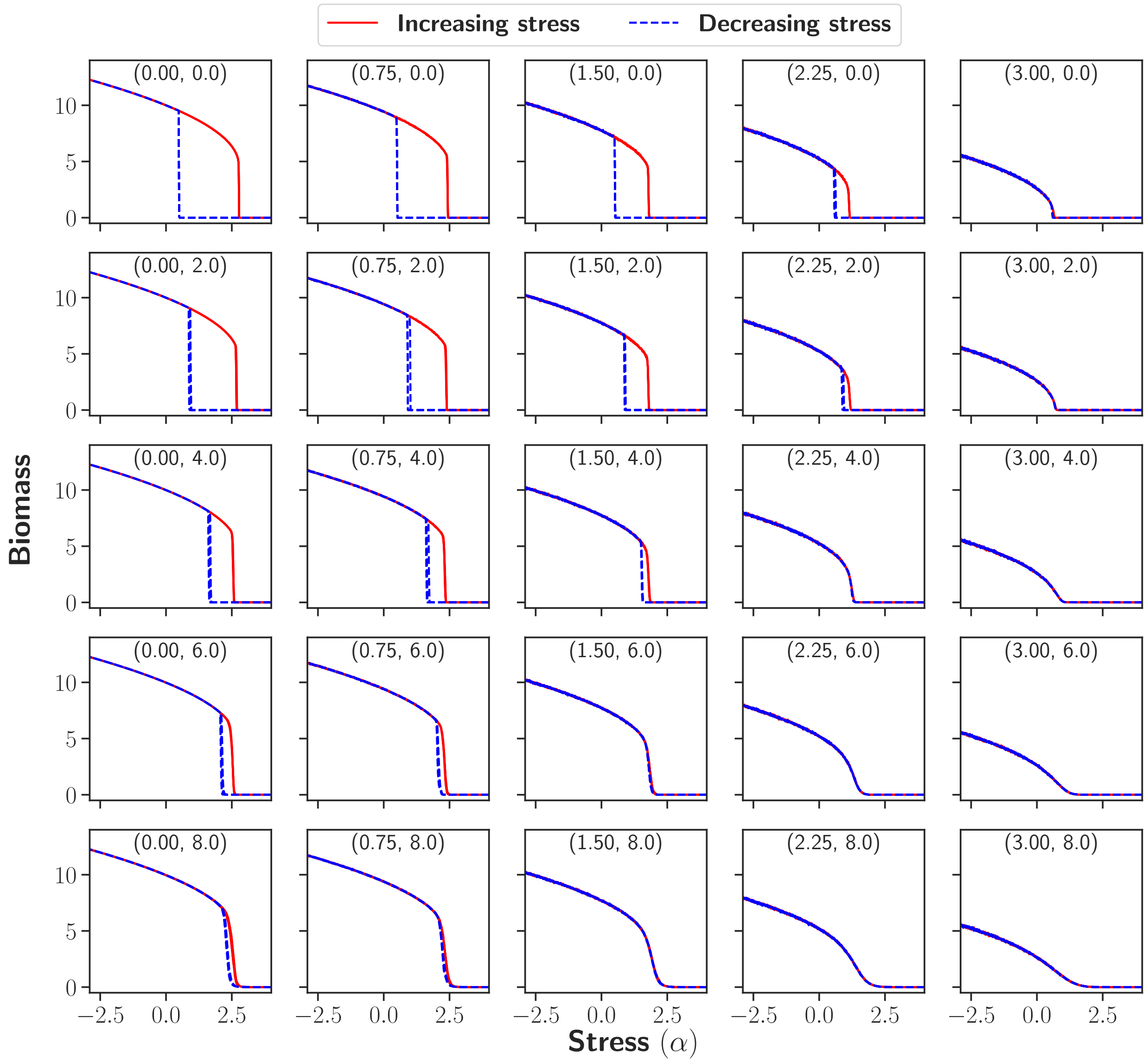


FIG. S79. Hysteresis diagrams showing the impact of various $(\sigma_e, \Delta)$ configurations in the grazing model. Model parameters: $r = 1$, $K = 10$, $\delta = 0.5$, dispersal rate $D = 1$, sweep rate $\epsilon = 10^{-5}$, and the system size $L \times L = 256^2$. The relevant value of $(\sigma_e, \Delta)$ appears at the top of each sub-panel.

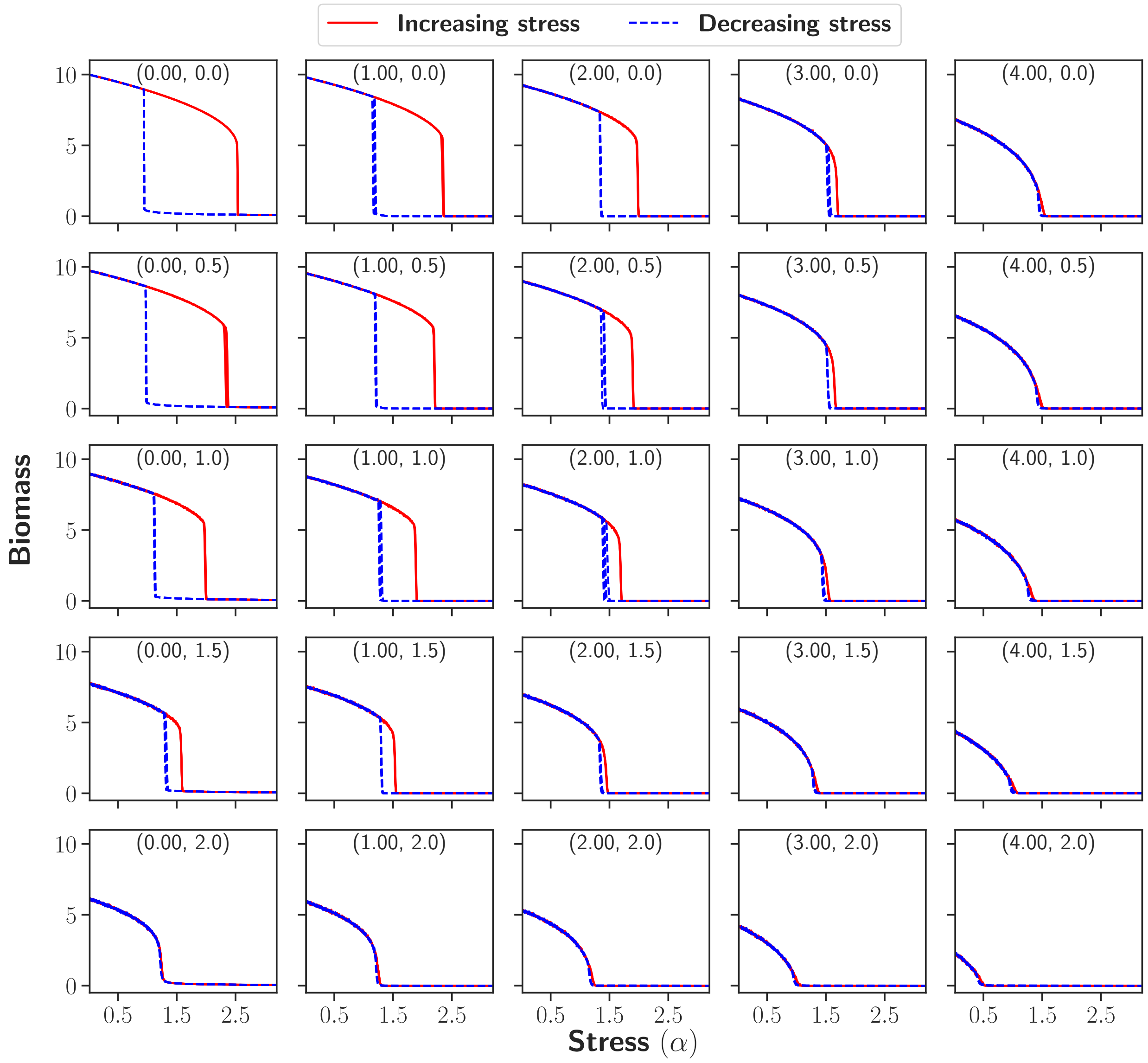


FIG. S710. Hysteresis diagrams illustrating the effects of different $(\sigma_d, \sigma_e)$ combinations in the insect outbreak model. Model parameters: $r = 1$, $K = 10$, $\delta = 0.5$, dispersal rate $D = 1$, sweep rate $\epsilon = 10^{-5}$, and the system size $L \times L = 256^2$. The relevant value of $(\sigma_d, \sigma_e)$ appears at the top of each sub-panel.

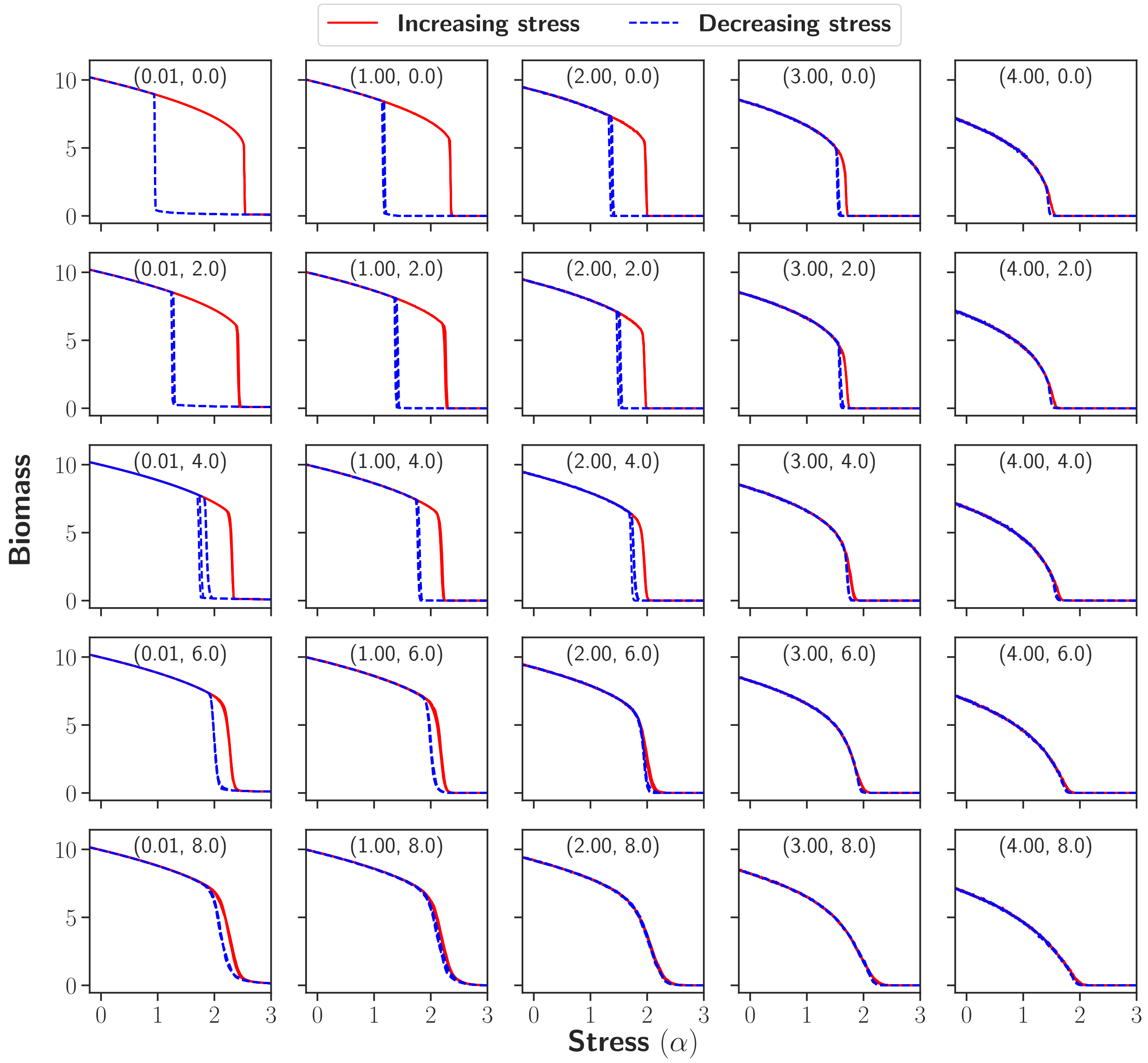


FIG. S711. Hysteresis diagrams showing the impact of various $(\sigma_d, \Delta)$ configurations in the insect outbreak model. Model parameters: $r = 1$, $K = 10$, $\delta = 0.5$, dispersal rate $D = 1$, sweep rate $\epsilon = 10^{-5}$, and the system size $L \times L = 256^2$. The relevant value of $(\sigma_d, \Delta)$ appears at the top of each sub-panel.

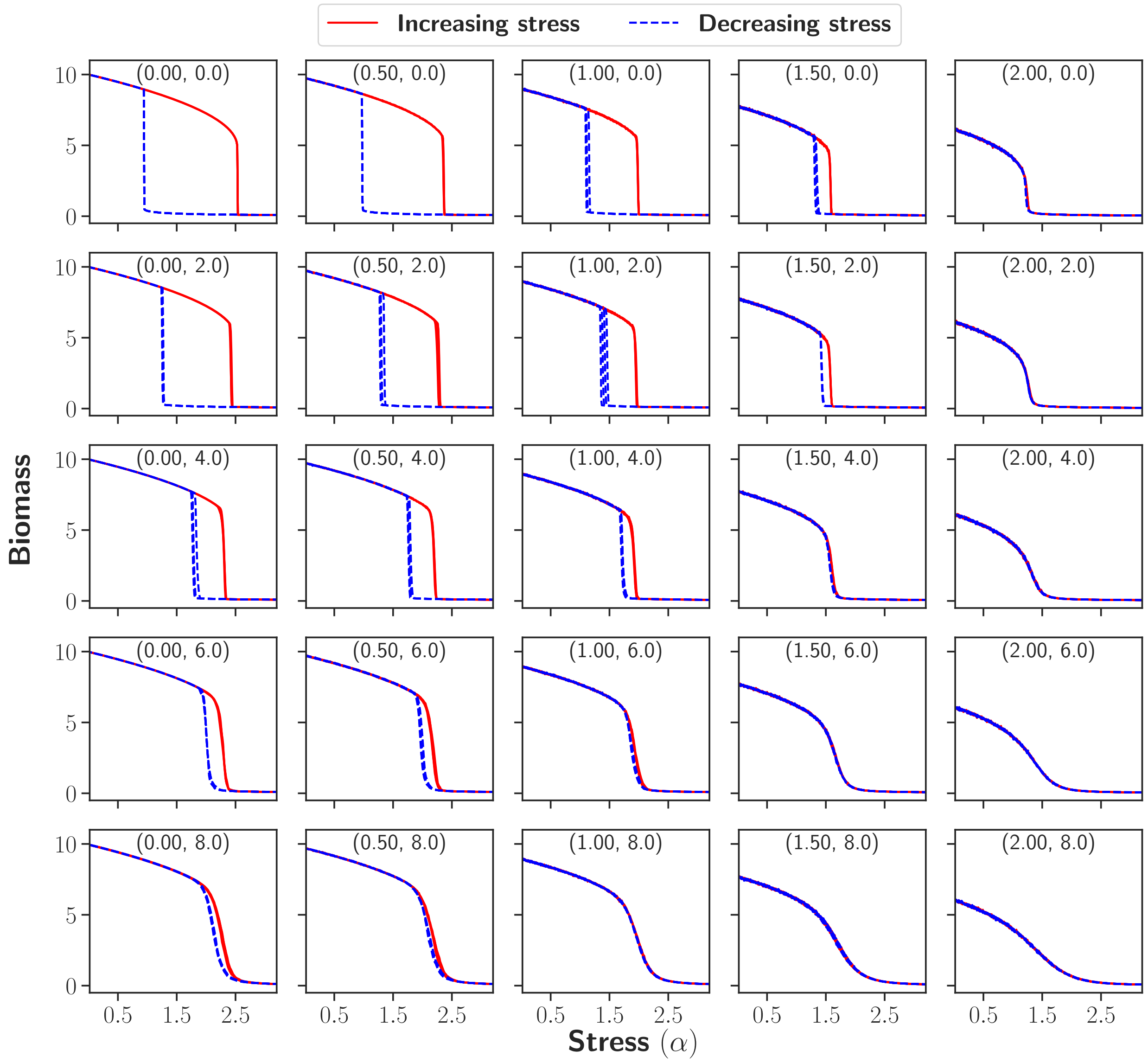


FIG. S712. Hysteresis diagrams representing the influence of changing $(\sigma_e, \Delta)$ values in the insect outbreak model. Model parameters: $r = 1$, $K = 10$, $\delta = 0.5$, dispersal rate $D = 1$, sweep rate $\epsilon = 10^{-5}$, and the system size $L \times L = 256^2$. The relevant value of $(\sigma_e, \Delta)$ appears at the top of each sub-panel.

##### T. Constructing heatmaps of DI and RGI in parameter planes - methodology

To construct heatmaps for a given parameter plane [e.g., $(\sigma_e, \Delta)$], we evaluate the chosen metric (e.g., DI) on a dense grid over an appropriate range and visualize the resulting field as a heatmap (see FIG. S713a). To improve accuracy, we use the average of four independent realizations and, for convenience, normalize each index to the interval $[0, 1]$. For clarity, we overlay contours (isoclines) at fixed levels 0.1–0.9 (see FIG. S713b). To obtain a smooth, gap-free surface and clearer isoclines, we apply cubic interpolation followed by Gaussian smoothing along both axes (see FIG. S713c).

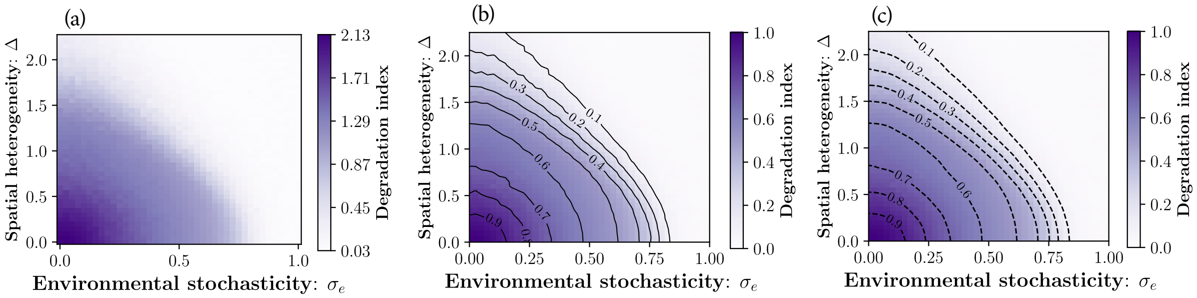


FIG. S713. Heatmaps of the degradation index (DI) in the $(\sigma_e, \Delta)$ plane for the lake-pollution model ($\beta = \delta = r = 1$, $p = 10$, $L^2 = 256^2$ with $\epsilon = 10^{-5}$). (a) A dense $40 \times 40$ grid over $\sigma_e \in [0, 1]$ and $\Delta \in [0, 2.25]$. (b) Isoclines at fixed levels 0.1–0.9, computed from the mean of four independent realizations and shown for the normalized DI (scaled to $[0, 1]$). (c) Cubic interpolation followed by Gaussian smoothing yields a gap-free surface with clearer isoclines.

##### U. Heatmaps of degradation and restoration-gap indices in the parameter planes

In what follows, we present heatmaps of the degradation index (DI) and the restoration-gap index (RGI) in the parameter planes $(\sigma_d, \sigma_e)$, $(\sigma_d, \Delta)$, and $(\sigma_e, \Delta)$ for three models, presented in sequence. (The corresponding results for the lake pollution model are shown in the main text and are therefore omitted here.)

Across all cases, the combined effect of the complexities, as reflected by the isoclines, is approximately additive. For example, in the desertification model (Fig. S714), the isoclines in the $(\sigma_d, \sigma_e)$ plane are parabolic and can be represented by an expression of the form $c_1\sigma_d^2 + c_2\sigma_e^2$, where $c_1$ and $c_2$ are constants determined by the model parameters. This additive behavior, consistent with a linear-response picture, breaks down when $\sigma_d$ or $\sigma_e$ become too large; however, in these regimes the associated regime shift is, in any case, much less "catastrophic."

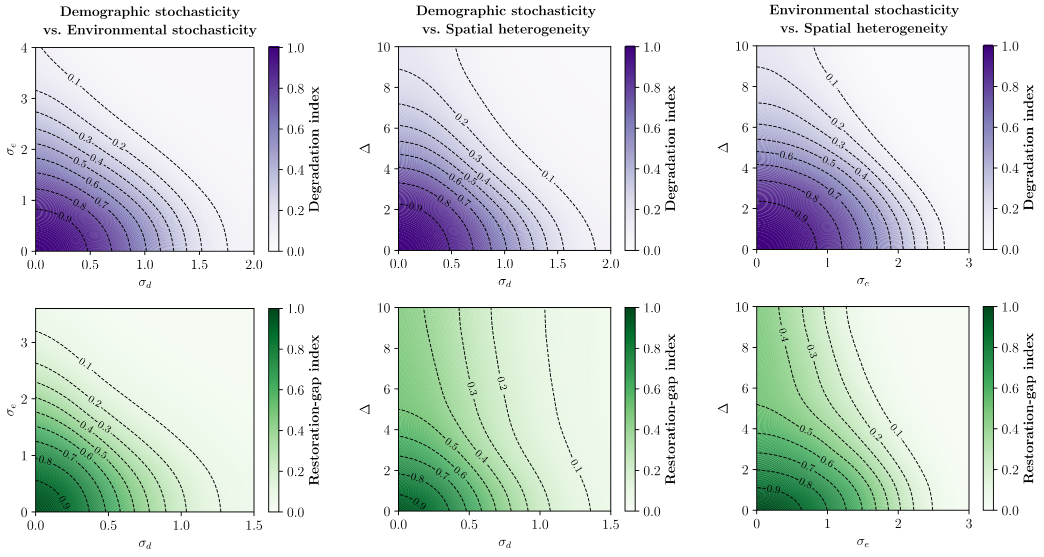


FIG. S714. **Desertification model:** Heatmaps of the normalized degradation and restoration–gap indices are shown in the first, and second rows, respectively, across the parameter planes $(\sigma_d, \sigma_e)$, $(\sigma_d, \Delta)$, and $(\sigma_e, \Delta)$ in the first, second, and third columns, respectively. Black dashed contours indicate isoclines at fixed levels from 0.1 to 0.9. In all cases we consider: $\beta = 2$, $\gamma = D = 1$, $L^2 = 256^2$, $\epsilon = 4 \times 10^{-5}$.

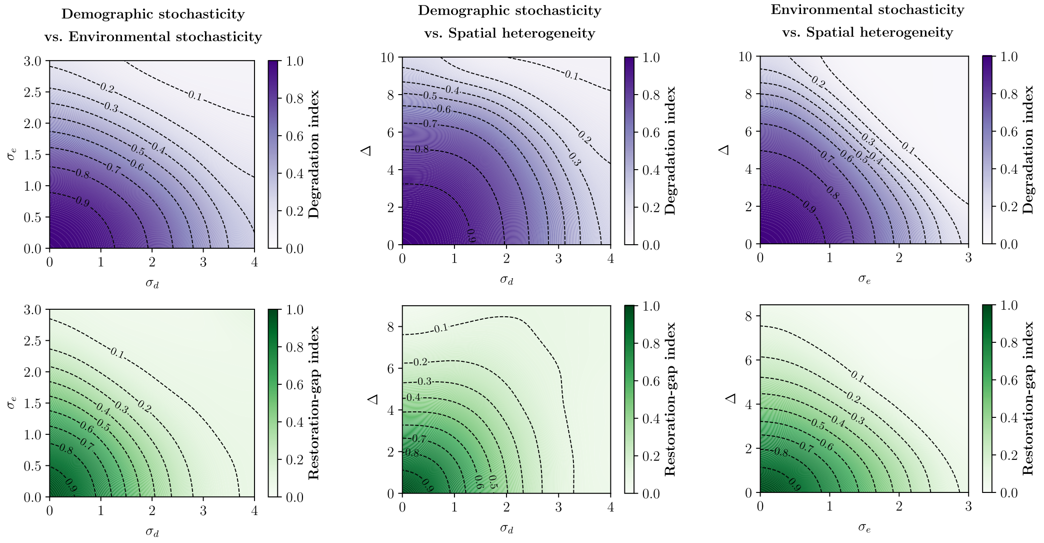


FIG. S715. **Grazing model:** Heatmaps of the normalized degradation and restoration–gap indices are shown in the first, and second rows, respectively, across the parameter planes $(\sigma_d, \sigma_e)$, $(\sigma_d, \Delta)$, and $(\sigma_e, \Delta)$ in the first, second, and third columns, respectively. Black dashed contours indicate isoclines at fixed levels from 0.1 to 0.9. In all cases we consider: $r = D = 1$, $K = 10$, $\delta = 0.5$, $L^2 = 256^2$, $\epsilon = 4 \times 10^{-5}$.

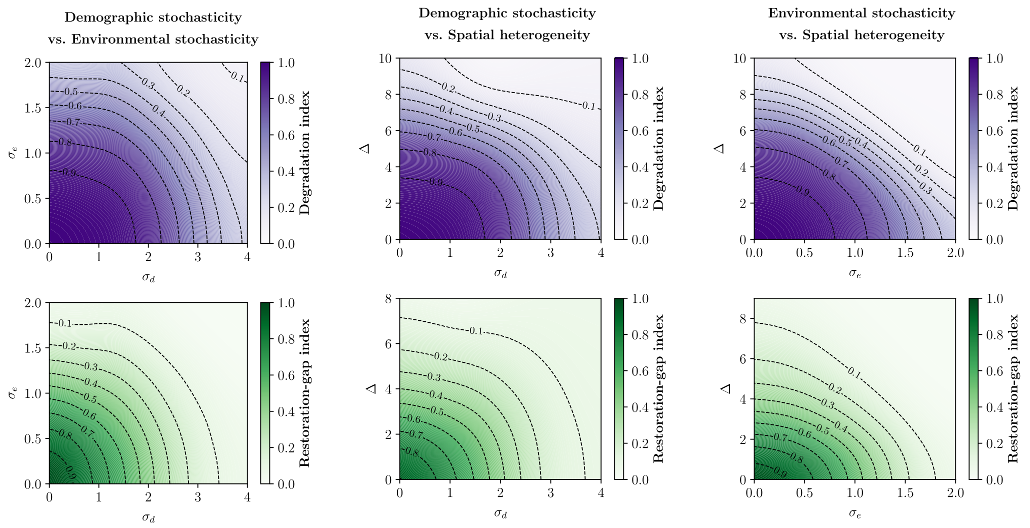


FIG. S716. **Insect outbreak model:** Heatmaps of the normalized degradation and restoration–gap indices are shown in the first, and second rows, respectively, across the parameter planes $(\sigma_d, \sigma_e)$, $(\sigma_d, \Delta)$, and $(\sigma_e, \Delta)$ in the first, second, and third columns, respectively. Black dashed contours indicate isoclines at fixed levels from 0.1 to 0.9. In all cases we consider: $r = D = 1$, $K = 10$, $\delta = 0.5$, $L^2 = 256^2$, $\epsilon = 4 \times 10^{-5}$.

### S8: Computational Methods implemented for Hysteresis loops and Phase Diagrams

In this section, we discuss in detail our numerical methodology used for constructing the phase diagrams. We first explain the parameter sweep approach (Subsection V) and then the marginal dynamics approach (Subsection W).

##### V. Parameter sweep approach: hysteresis diagram

To construct a full hysteresis diagram, we employ a parameter-sweep method in which all parameters are fixed except the stress parameter $\alpha$. We then perform an upward sweep, in which $\alpha$ is gradually increased in time at a prescribed rate while tracking the biomass density, with the system initialized in a healthy state so that, as stress increases, it deteriorates to a degraded condition. This is followed by a downward sweep, in which $\alpha$ is gradually decreased in time while tracking the biomass density, using the final state of the system at the end of the upward sweep as the initial condition, allowing the system to recover back to the healthy state. The (history dependent) state of the system through upward and downward sweeps, as a function of stress $\alpha$, yield the complete hysteresis diagram.

To allow the system to relax toward equilibrium, we consider slow sweep rates. Specifically, we use $\alpha(t) = \alpha_{\min} + \epsilon t$ for the upward sweep and $\alpha(t) = \alpha_{\max} - \epsilon t$ for the downward sweep, where $\alpha_{\max}$ and $\alpha_{\min}$ define the window over which $\alpha$ is varied. Throughout the paper, we use sweep rate $\epsilon = 10^{-4}$, $4 \times 10^{-5}$, and $10^{-5}$.

It is worth mentioning that in systems with an absorbing state, such as the desertification and grazing models, the system becomes trapped in the absorbing state with zero biomass density at the end of the upward sweep. When this state is used as the initial condition for the downward sweep, the system cannot recover to the healthy state. To overcome this, we introduce a minute amount of external migration $\mu$ at each lattice site after each time interval. Throughout the paper, we consider $\mu = 10^{-5}$ and $10^{-4}$.

##### W. Marginal dynamics identification

The parameter sweep method described in the last section is used to identify the collapse and the recovery tipping points. In parallel with this method we have implemented the marginal dynamics approach described below. For the tipping points, this approach serves as a consistency check on the first: since both methods compute the same quantity, their results should coincide. Moreover, the marginal dynamics approach allows us to identify the Maxwell point and the continuum transition point (when exists).

In this Subsection we present the computational details of four different types of transition points mentioned through the main text.

**Collapse tipping point:** At the tipping point, one of the stable states of the system loses its stability. When the system is trapped in a stable or metastable state, it crosses over to the alternative state only due to the effect of stochasticity, therefore the process is slow, and becomes slower as stochasticity decreases. On the other hand, once the state losses its stability, the deterministic dynamics takes the system to the alternative state so the process is quite fast.

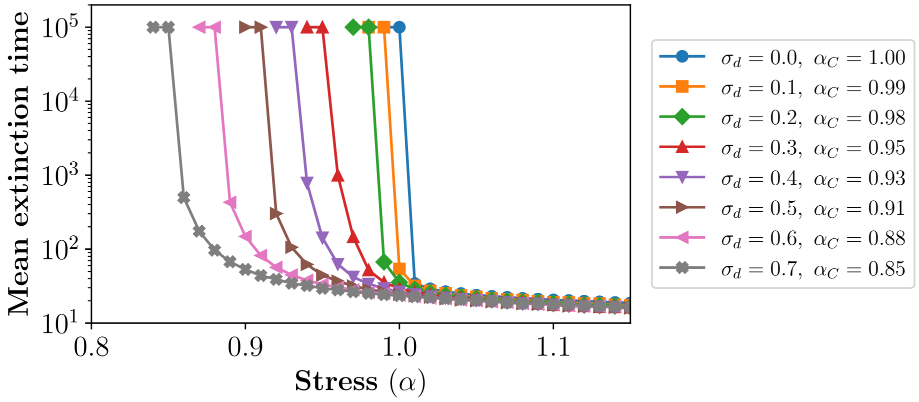


FIG. S81. Mean extinction time as a function of the stress parameter $\alpha$, shown for different values of $\sigma_d$, in the desertification model with $\beta = 2$, $\gamma = D = 1$, and $L^2 = 256^2$. Curves are shown for eight $\sigma_d$ values, and each point is an average over 32 realizations. For each $\sigma_d$, the corresponding $\alpha_C$ is defined as the smallest $\alpha$ plotted for that case minus the step size between consecutive $\alpha$ values (0.01).

To identify the collapse tipping point ($\alpha_C$), the system is initialized in a healthy state (e.g., vegetation in the desertification model) and its mean extinction time is monitored for a given and fixed stress parameter $\alpha$. As $\alpha$ is decreased from a high value ($\alpha > \alpha_C$), the mean extinction time rises gradually. Close to the tipping point, for values just above $\alpha_C$, the mean extinction time is estimated by averaging over many realizations. Below $\alpha_C$, the healthy state becomes stable and extinction times grow extremely large, making direct estimation impractical within feasible simulation times. An upper cutoff is therefore imposed on the simulation runtime; if the measured mean extinction time reaches this cutoff, the corresponding $\alpha$ value is treated as lying at or just below the tipping point (see FIG. S81).

Alternatively, we can consider a system initially in a healthy state and examine, again at fixed $\alpha$, its time evolution over a suitable range of the stress parameter $\alpha$. When $\alpha$ is below the tipping point, the system remains in the healthy state even at long times. However, above the tipping point, the system makes a transition to a degraded state (FIG. S82(a)). Therefore, if we plot the biomass density (measured at large times) against $\alpha$, the abrupt drop in biomass density from the healthy to the degraded state allows us to identify the collapsed tipping point (FIG. S82(b)).

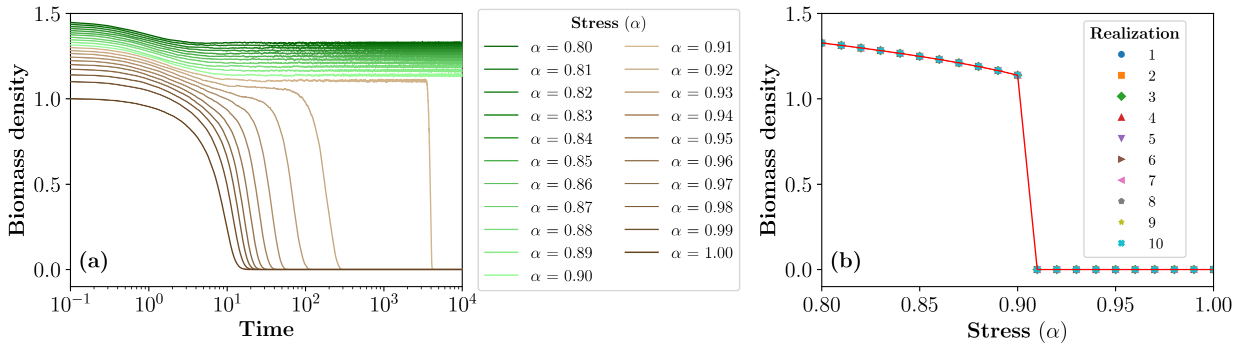


FIG. S82. Desertification model with $\beta = 2$, $\gamma = D = 1$, $\sigma_d = 0.5$, and $L \times L = 256^2$. (a) Biomass density plotted versus time for selected values of stress parameter $\alpha$. (b) Biomass density at late time ($t = 10000$) plotted against $\alpha$, displaying a sharp transition at $\alpha = 0.90$. For each $\alpha$, 10 points are shown, which fall on top of each other, each corresponding to an independent realization.

Notably, this procedure is valid for all models. For example, in the lake pollution model, the degraded state, characterized by turbid water, corresponds to a non-zero solution. Unlike in the desertification model, this makes it difficult to define an extinction time for the system. However, in the same system, by examining the time evolution of water quality (defined as the inverse of biomass density) and measuring it at long times, one can observe an abrupt shift in water quality (see FIG. S83).

**Recovery Tipping Point:** Here we explain the methods by which we identify the recovery tipping point. We first discuss models with an absorbing state (e.g., the desertification model) and then models without an absorbing state (e.g., the lake pollution model).

First consider the model with an absorbing state. Specifically, for the desertification model, the system is initialized in a vegetated state with a very low biomass density. $\alpha$ is kept fixed through the numerical experiments, and we identify the tipping point by examining the dynamics at different values of $\alpha$.

At the recovery tipping point, the bare-soil state us marginally stable, therefore one expects a

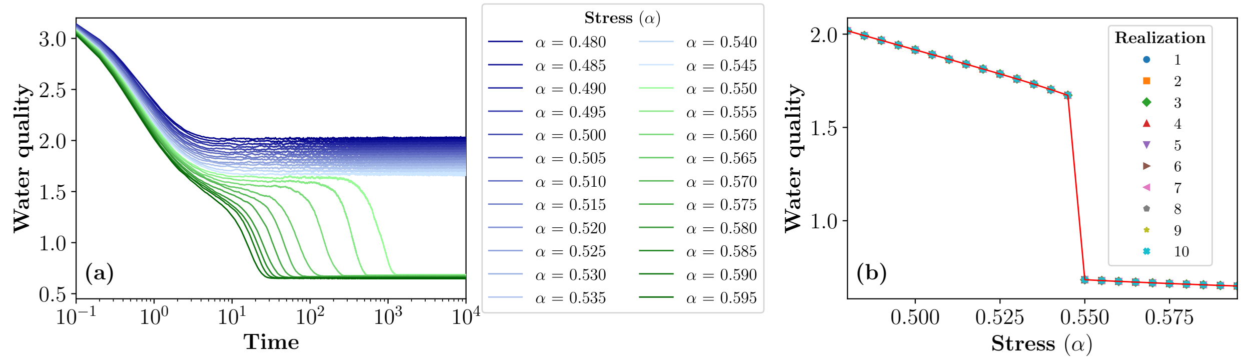


FIG. S83. Lake pollution model with $\beta = r = \delta = D = 1$, $p = 10$, $\sigma_d = 0.5$ and $L \times L = 256^2$. (a) Water quality, defined as the inverse of biomass density, plotted versus time for selected values of stress parameter $\alpha$. (b) Water quality recorded at large times ($t = 10000$) plotted versus $\alpha$, indicating a sudden jump at $\alpha = 0.545$. For each $\alpha$, 10 points are shown, which fall on top of each other, each corresponding to an independent realization.

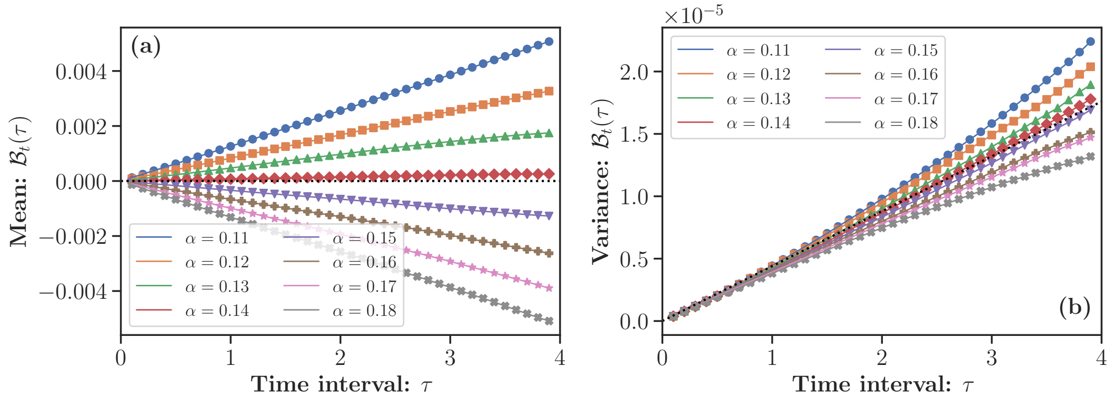


FIG. S84. Desertification model with parameters: $D = 1$, $\sigma_d = 0.5$, $\beta = 2$, $\gamma = 1$, and lattice size $L^2 = 256^2$. The mean (a) and variance (b) of the quantity $\mathcal{B}_t(\tau)$ are plotted as functions of the time interval $\tau$, for the same eight values of $\alpha$. For $\alpha = 0.14$, the mean remains close to zero across all $\tau$, while the variance increases approximately linearly. This behavior is consistent with neutral dynamics and indicates that the recovery tipping point for this set of parameters occurs at $\alpha = 0.14$. (A total of 2400 independent realizations were conducted to produce the results).

neutral dynamics at its vicinity. Under neutral dynamics, the quantity [27],

$$\mathcal{B}_t(\tau) = \frac{B(t+\tau) - B(t)}{\sqrt{B(t)}}$$

is a process with zero mean, and its variance grows linearly with the time interval *tau*. This behavior allows us to pinpoint the value of $\alpha$ corresponding to the recovery tipping point (see FIG. S84).

Using the same initialization, we estimate the recovery tipping point through an alternative approach. In the deterministic model, the recovery threshold is known to occur for the desertification model at $\alpha = 0$. However, when demographic stochasticity is present, this threshold effectively shifts, and the renormalized stress parameter $\alpha_R$ reaches zero at the tipping point. To estimate $\alpha_R$, we compute the finite-difference approximation of the growth rate,

$$\frac{\Delta B}{\Delta t} = \frac{B(t + \Delta t) - B(t)}{\Delta t},$$

and plot it against $B(t)$ for a given $\alpha$, where $\Delta t$ is a small time interval. By aggregating a large number of such $(B(t), \text{growth rate})$ pairs and performing an ordinary least squares linear regression, we obtain a best-fit line whose slope provides an estimate of $\alpha_R$. Repeating this procedure across a range of $\alpha$ values allows us to identify the value for which the slope is closest to zero, thereby determining the recovery tipping point in the presence of demographic stochasticity (see FIG. S85).

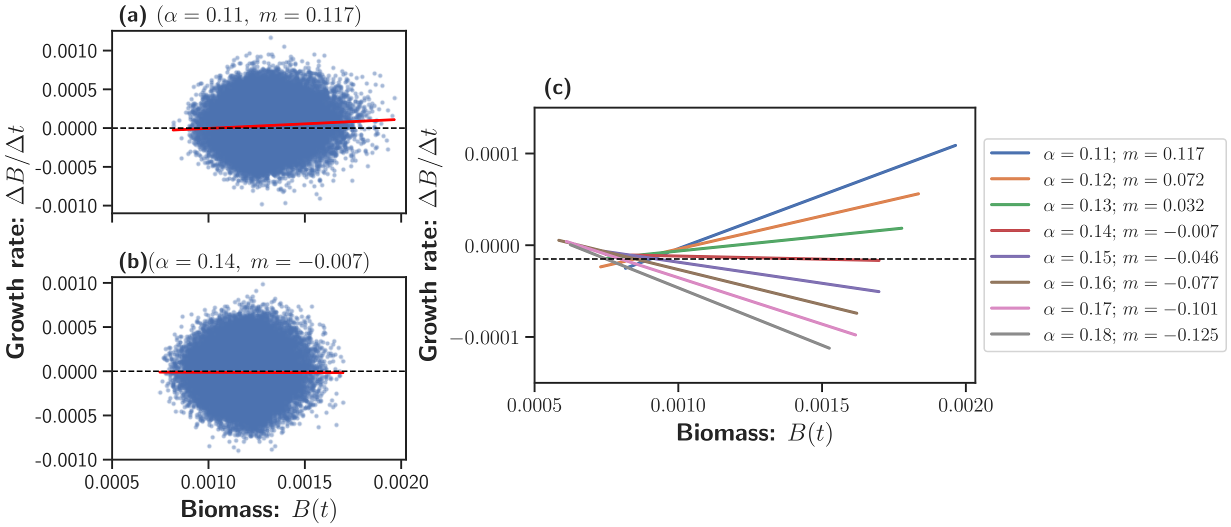


FIG. S85. Desertification model with parameters: $D = 1$, $\sigma_d = 0.5$, $\beta = 2$, $\gamma = 1$, and $L^2 = 256^2$. The initial population at each site was drawn from a Gaussian distribution with mean zero and variance $10^{-5}$; negative values were set to zero. A total of 2400 independent realizations were used to generate the results shown above. In (a) and (b) $\Delta B/\Delta t$ is plotted against $B(t)$ for two different values of $\alpha$. Each blue point represents a single data point, computed for $t \leq 4$ with $\Delta t = 0.1$. For each $\alpha$, a best-fit line (shown in red) is obtained using ordinary least squares regression. (c) The best-fit lines for all $\alpha$ values are plotted together to identify the tipping point based on the slope $(m)$. For this set of parameters, the recovery tipping point occurs at $\alpha = 0.14$, as the corresponding slope is closest to zero.

For the model without an absorbing state, such as the lake pollution model, the recovery tipping point is identified using a procedure analogous to that employed earlier to determine the collapse tipping point in the same system. Specifically, we initialize the system in the degraded state and track its time evolution across a suitable range of the stress parameter $\alpha$. When $\alpha$ is above the tipping point, the system remains in the degraded state even at long times. In contrast, at or below the tipping point, the system makes an abrupt jump to the healthy state (see in FIG. S86(a)). By plotting the water quality at large times as a function of $\alpha$, the tipping point can be identified (see FIG. S86(b)).

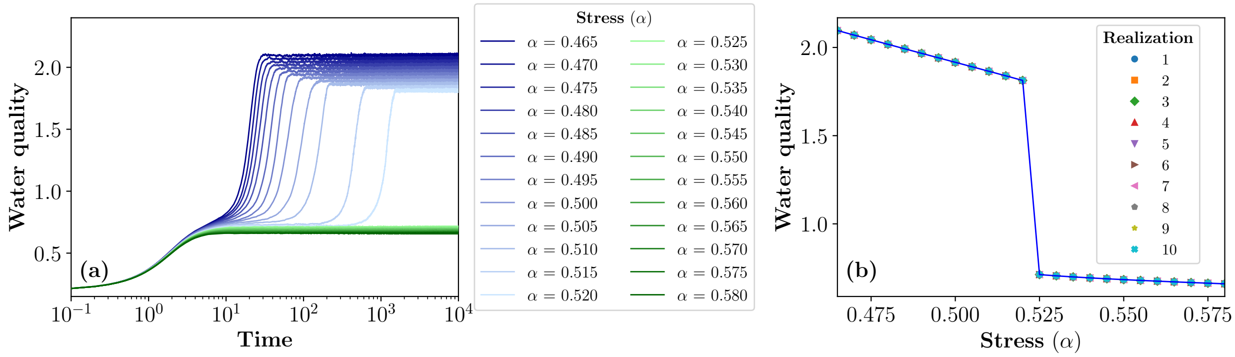


FIG. S86. Lake pollution model with $\beta = r = \delta = D = 1$, $p = 10$, $\sigma_d = 0.5$, and $L \times L = 256^2$. (a) Time series of water quality (defined as the inverse of biomass density) for various values of the stress parameter $\alpha$. (b) Water quality measured at large time ($t = 10000$) is plotted as a function of $\alpha$, revealing an abrupt transition near $\alpha = 0.520$. For each $\alpha$, 10 points are shown, which fall on top of each other, each corresponding to an independent realization.

**Maxwell Point:** The front velocity between healthy (e.g., vegetation) and degraded (e.g., bare-soil) states vanishes at the Maxwell point [28]; consequently, after an initial transient, the average biomass density becomes time invariant. Figures S87(a) and S87(b) show, respectively, six snapshots of the system and the corresponding biomass-density trajectories for three values of the stress parameter $\alpha$ that lie just below, near, and just above the Maxwell point.

When $\alpha$ is below the Maxwell point, vegetation gradually invades bare soil (first row in Fig. S87(a)), and the late-time biomass density increases (blue curve in Fig. S87(b)). When $\alpha$ is above the Maxwell point, bare soil invades vegetation (third row in Fig. S87(a)), leading to a decline in biomass density (green curve in Fig. S87(b)). Near the Maxwell point, the phase boundary is stationary (second row), as the invasion velocity vanishes, and the late-time biomass density remains constant (orange curve).

**Continuous transition to an absorbing state:** As discussed in the main text, the biomass

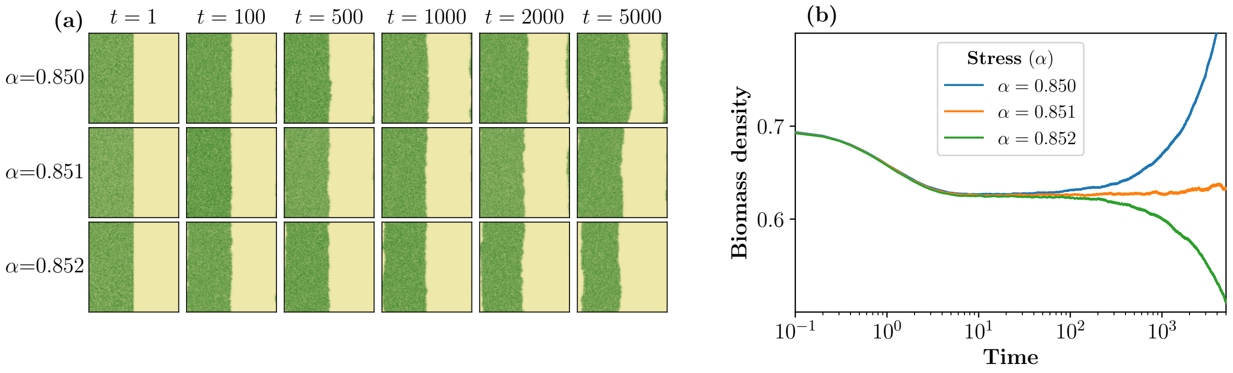


FIG. S87. Desertification model with $\beta = 2$, $\gamma = D = 1$, with $\sigma_d = 0.5$, and $L \times L = 256^2$. (a) Snapshots at six times for three values of the stress parameter $\alpha$, just below, at, and just above the Maxwell point (top to bottom). (b) Corresponding biomass density versus time, averaged over 24 realizations; the orange curve denotes the Maxwell-point case.

density undergoes a continuous transition in the limit of high stochasticity in models that possess an absorbing state (e.g., the desertification and grazing models). Villa *et al.* [2] pointed out that, in the presence of an absorbing state, increasing demographic stochasticity drives the system from a hysteretic, discontinuous transition to a continuous, non-hysteretic one. It was also shown that this transition belongs to the directed-percolation universality class [10, 11].

To identify the transition point, we rely on the known critical exponents of the directed-percolation transition. For a given stress parameter $\alpha$, the system is initialized in the healthy state and let evolve in time, and the mean biomass density is monitored through the process.

The possible outcomes are illustrates in Figure S88(a). After an initial transient time, for $\alpha < \alpha_C$ the total biomass density eventually saturates to a finite value, whereas for $\alpha > \alpha_C$, it decays exponentially at late times and eventually reaches the absorbing state. Near the critical point, $\alpha \simeq \alpha_C$, the density exhibits a power-law decay in time, characterized by the critical exponent $\bar{\delta} \simeq 0.45$ (Fig. S88(b)), consistent with density-decay exponent of the directed-percolation universality class [10].

Figure S88(c) shows the steady state biomass density (when demographic stochasticity is large) as a function of the stress parameter. This density decreases continuously with $\alpha$ until it reaches zero at some critical value $\alpha_c$. By replotting the density as a function of the rescaled parameter $(\alpha_c - \alpha)$, we verify that the mean biomass density follows the predicted power-law scaling with exponent $\bar{\beta}_{2d} \simeq 0.58$ (Fig. S88(d)), consistent, again, with order parameter exponent of the directed-percolation universality class [10].

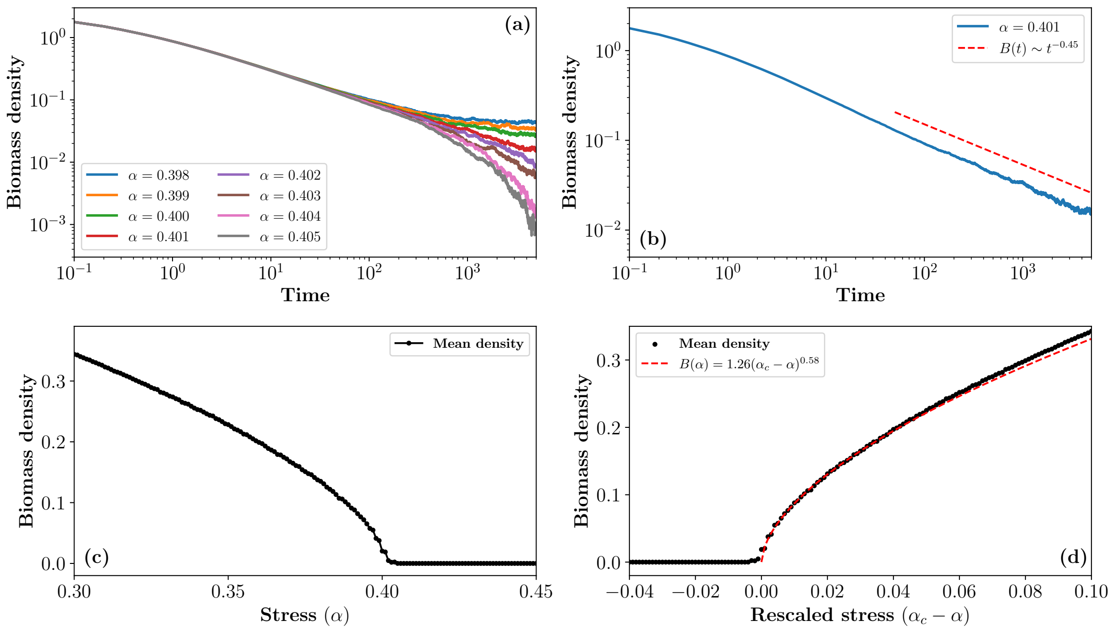


FIG. S88. Desertification model results are shown for parameters $\beta = 2$, $\gamma = D = 1$, with $\sigma_d = 2.0$ and $L \times L = 256^2$. (a) Time evolution of the mean biomass density is shown for a range of $\alpha$ values, illustrating the transition dynamics. (b) At the critical stress level $\alpha \simeq \alpha_c$, the time evolution of biomass density exhibits a power-law decay, characterized by the exponent $\bar{\delta} \simeq 0.45$. Each data point in the plots represents an average over 8 independent realizations. (c) The late-time ($t > 4000$) mean biomass density is plotted as a function of the stress parameter $\alpha$, with the system initialized in the healthy (e.g., vegetation) state corresponding to the deterministic model. (d) The same mean biomass density is plotted against the rescaled stress parameter $(\alpha_c - \alpha)$, demonstrating a power-law scaling behavior characterized by the exponent $\bar{\beta}_{2d} \simeq 0.58$.

---

[7] R. Lande, S. Engen, and B.-E. Saether, *Stochastic population dynamics in ecology and conservation* (Oxford University Press, USA, 2003).

[20] M. Scheffer, *Critical transitions in nature and society* (Princeton university press, 2009).
